# STcompare: comparative spatial transcriptomics data analysis of structurally matched tissues to characterize differentially spatially patterned genes

**DOI:** 10.1101/2025.11.21.689847

**Authors:** Kalen Clifton, Vivien Jiang, Rafael dos Santos Peixoto, Srujan Singh, Ryo Matsuura, Hamid Rabb, Jean Fan

## Abstract

**Motivation:** Comparative analysis of spatial transcriptomics (ST) data is needed to identify genes that spatially change in their expression patterns between conditions, such as in diseased versus healthy tissues. Existing methods generally fail to distinguish changes in spatial patterning by focusing only on changes in gene expression magnitude for methods adapted from non-spatial data or on changes in significance of spatial variability for methods focusing on spatially-resolved data.

**Results:** To address these limitations, we develop STcompare, a statistical framework for comparative analysis of ST data by testing for differences in spatial correlation and spatial fold-change across structurally matched locations. Using simulated data, we demonstrate how STcompare provides distinct insights from bulk differential gene expression analysis and spatially variable gene expression analysis as well as other spatial comparison methods. STcompare further robustly controls for false positives even in the presence of spatial autocorrelation common in ST data. We apply STcompare to real ST data of biological replicates of mouse brains to confirm high spatial correspondence of gene expression patterns across samples. We apply STcompare to identify genes that spatially change in mouse kidneys with acute kidney injury compared to a healthy control, revealing tissue compartment-specific molecular dysregulation. Overall, the application of this spatially-aware comparative analysis will enable the discovery of differential spatially patterned genes across various physiological and technological axes of interest.

**Availability and Implementation:** STcompare is implemented as an open-source R package at https://github.com/JEFworks-Lab/STcompare with additional documentation and tutorials available at https://jef.works/STcompare/.

## Introduction

Spatial transcriptomic (ST) technologies enable the investigation of how tissue organization relates to cellular function by capturing the spatial location of cells within a tissue and the gene expression profiles associated with those locations^1–3^. As ST technologies are increasingly applied in the context of disease characterization and translational research, identifying genes that spatially change in their expression patterns in diseased tissues as compared to healthy controls can reveal localized molecular variation associated with pathological processes, offering insights into disease mechanisms as well as aiding in the discovery of therapeutics and diagnostic biomarkers^4,5^.

Attempts at such comparative analysis of ST datasets can be performed using traditional non-spatial bulk or single cell transcriptomics analysis approaches such as differential gene expression (DGE) analysis, comparing the expression magnitudes of genes between two datasets. However, this type of DGE analysis does not consider the spatial information. As such, a gene can be identified as not differentially expressed across two datasets by having the same mean gene expression despite having distinct spatial expression patterns. Conversely, a gene can be identified as differentially expressed across the two datasets by having significantly different mean gene expression despite having spatial expression patterns that are highly similar. Likewise, spatially variable gene (SVG) expression analysis can distinguish spatially variable from non-spatially variable genes across conditions. But in a comparative setting, a gene can be identified as spatially variable despite having distinct spatial expression patterns. As such, existing DGE and SVG analyses fail to characterize changes in spatial expression patterns.

To demonstrate these cases, we simulated ST datasets where one pair of genes, A and B, had different radial expression patterns but the same mean expression, while the other pair, A and C, had same radial expression patterns but different mean expression (Figure 1a). Indeed, when we applied non-spatially resolved bulk DGE analysis, the former spatially different pair was classified as not significantly differentially expressed (log_2_ fold-change = −0.03, Wilcoxon rank sum test p-value = 0.39) and the latter similarly patterned pair was classified as significantly differentially expressed (log_2_ fold-change = 1.61, Wilcoxon rank sum test p-value < 2.2e-16). Likewise, when we applied SVG analysis via Moran’s *I* genes A, B, and C are all identified as significant spatially variable genes (A *I* = 0.82 *p* < 1e-16; B *I* = 0.83 *p* < 1e-16; C *I* = 0.80 *p* < 1e-16) such that significance cannot be used to distinguish which genes are spatially different from each other (Supplementary Methods). These results demonstrated that bulk DGE and SVG analysis results do not reflect changes or lack of changes in spatial expression patterns.

**Figure 1.**
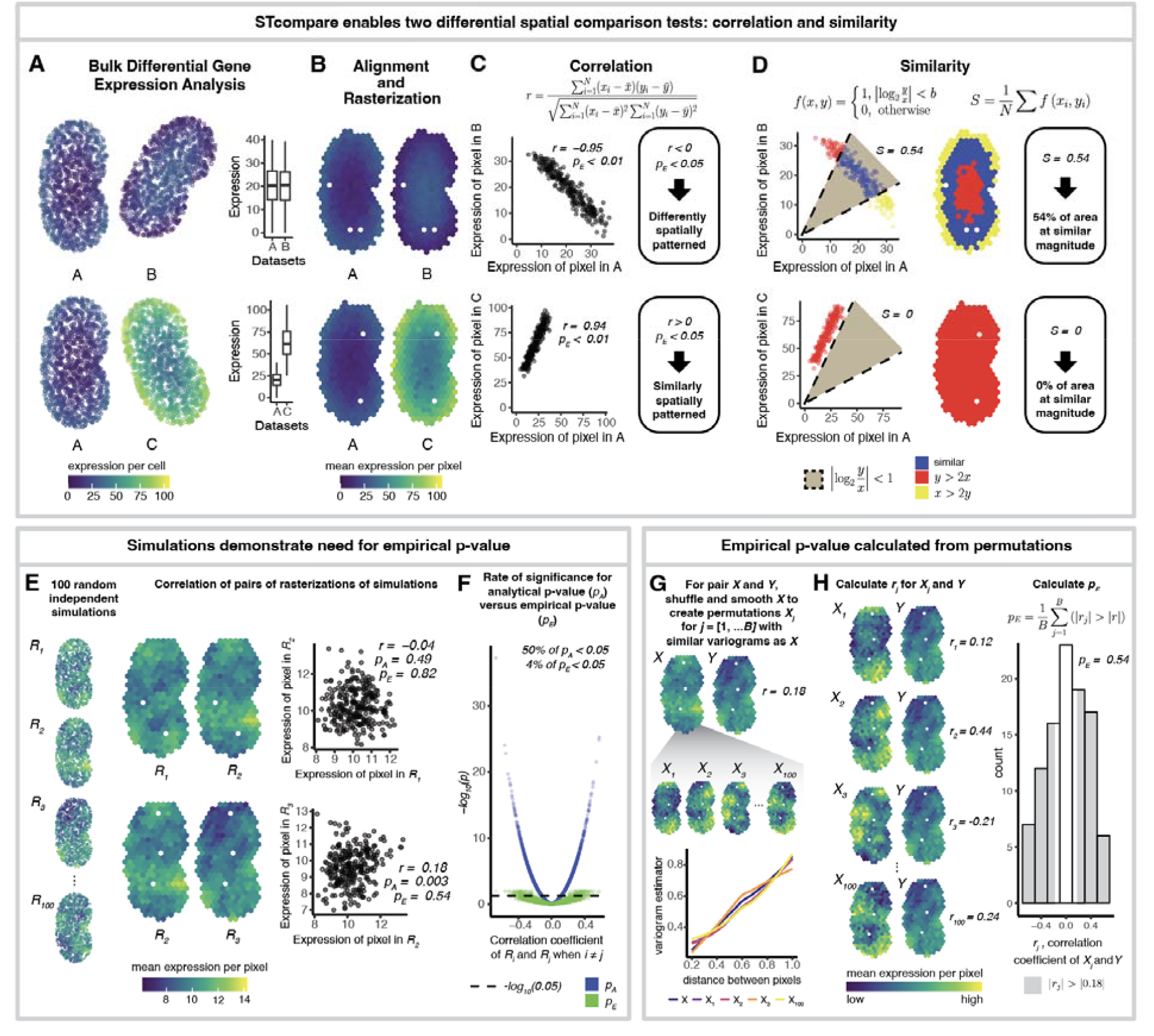
STcompare enables differential spatial comparison tests distinct from bulk differential gene (DGE) analysis. **A.** Simulated spatial transcriptomics data of a gene across three samples (A, B, and C). With bulk DGE methods, the gene is identified as not differentially expressed between samples A and B because they have the same mean gene expression even though they have different radial expression patterns (top). Conversely, the gene is identified as differentially expressed between samples A and C because they have very different mean gene expression even though they have the same radial expression patterns (bottom). **B**. Simulated spatial transcriptomics data after spatial structural alignment and rasterization onto a shared pixel grid to obtain matched spatial locations for evaluation with STcompare. **C**. Using STcompare’s spatial correlation test, gene expression patterns A and B are classified as differently spatially patterned (top) and gene expression patterns A and C are classified as similarly spatially patterned (bottom) since the pairs have statistically significant negative and positive empirical Pearson’s correlation coefficients, *r*, respectively. **D**. Using STcompare’s spatial fold-change test to calculate a similarity score, *S*, that gives the percentage of pixels that have an absolute log_2_ fold change < 1, 54% of the area of gene expression patterns A and B is expressed at similar magnitude (top) and 0% of the area of gene expression patterns A and C is expressed at similar magnitude (bottom). **E**. For the spatial correlation test, to demonstrate the need for empirical p-values, 100 genes with independent, random, autocorrelated patterns are simulated. Select simulated genes with rasterized outputs and correlation plots shown **F**. For all possible pairs (excluding self-pairs) amongst these 100 random genes, Pearson’s correlation coefficients are plotted versus the −log_10_ (*p*) for analytical (blue) and empirical (green) p-values. Using significance threshold of p-value > 0.05 (dashed line), 50% of these analytical p-values are significant, while only 4% of the empirical p-values are significant. **G** To calculate an empirical p-value for a pair of gene expression patterns *X* and *Y*, we create permutations, *X*_*j*_ for[1, …, *B*], where 100. Select permutations *X*_*1*_, *X*_*2*_, *X*_*3*_, and are shown (top). Permutations are created by shuffling and then smoothing so that the resulting permutations *X* _*j*_ and have similar variogram plots, a measure of the degree of autocorrelation. The variogram plot (bottom), which plots the variogram estimator as a function of distance between spatial locations, shows the values for (blue) and select permutations (purple), (pink), (orange) and (yellow) for distances within the 25^th^ percentile. **H**. Computing the correlation coefficients *r*_*j*_, for *Y* and each *X*_*j*_ for *j*[1, …, *B*] results in an empirical null distribution shown as a histogram of all correlation coefficients *r*_*j*_ (right). For select iterations, the coefficients and the rasterized gene expression of the pairs are shown (left). A two tailed empirical p-value, *p*_*E*_, is calculated as the proportion of *r*_*j*_ that have an absolute value greater than the absolute value of r, the correlation coefficient *X* for and *Y* (shaded region).

## Methods

To enable comparative analysis for ST datasets that considers spatial information in structurally matched tissues, we developed STcompare, available as an R package at https://github.com/JEFworks-Lab/STcompare. STcompare enables two differential spatial comparison tests: one based on spatial correlation, and one based on spatial fold change (Figure 1b-d). To use STcompare, ST datasets should be preprocessed to ensure matched spatial locations for comparison such as through alignment of shared tissue structures using previously developed approaches^6–12^ and rasterization^13^ onto a shared pixel grid as needed (Figure 1b).

The spatial correlation test of STcompare identifies gene expression patterns that have shifted in spatial dependencies across datasets. We have defined differently spatially patterned genes as those with statistically significant negative Pearson’s correlation coefficients and similarly spatially patterned as genes as those with statistically significant positive Pearson’s correlation coefficients (Figure 1c). Notably though, it has been described that using standard statistical tests, such as t-tests or Z-tests, to evaluate the Pearson’s correlation coefficient significance for spatially autocorrelated data results in a high false positive rate (FPR) because spatial autocorrelation violates the assumption of the parametric statistical tests that values are independently distributed^14–16^. Since spatially patterned genes have spatial autocorrelation^15,17–21^, we expected analytical p-values from such statistical tests would result in inflated number of genes identified as significantly correlated. To demonstrate this high FPR, we simulated 100 genes, each with an independent, random, autocorrelated pattern from a Gaussian random field and calculated the Pearson’s correlation coefficients and analytical p-values *p*_*A*_ for all pairs using a standard Z-test, which assumes no autocorrelation (Figure 1e, Supplementary Methods). We observed a high FPR of 0.50 with 4928 of 9900 pairs (excluding self-pairs) identified as significantly correlated using level of significance as p-value < 0.05 well beyond the expected FPR of 0.05 (Figure 1f). To address this problem, STcompare generates empirical null distributions that model spatial autocorrelation to calculate an empirical p-value using an approach previously described in Viladomat et al^14^ (Supplementary Methods). Briefly, given two datasets *X* and *Y*, STcompare creates permutations of *X* defined as *X*_*j*_ for *j* [1, …, *B*], which have the same degree of autocorrelation as the original *X*. Permutations are created by shuffling and then smoothing *X* with a Gaussian kernel to add back autocorrelation. To determine size of the kernel, we define *δ* as the percentage of neighbors that should be within the area enclosed by the radius of the Gaussian kernel. So that the permutations *X*_*j*_ have the same degree of autocorrelation as the original *X*, we choose the δ that minimizes the sum of squared errors between the variograms of *X* and each *X*_*j*_, since variograms are a function that plots the difference between points against the distance between them (Figure 1g). Next, computing the correlation coefficients *r*_*j*_ for *Y* and each *X*_*j*_ for *j* [1, …, *B*] results in an empirical null distribution. Finally, a two tailed empirical p-value is calculated as the proportion of correlation coefficients *r*_*j*_for *Y* and each *X*_*j*_ that have an absolute value greater than the absolute value of the correlation coefficient *r* for *X* and *Y* (Figure 1h, Supplementary Methods). When applied to the simulated data, this method of generating empirical p-values for Pearson’s correlation coefficients reduces the inflated number of pairs with autocorrelation identified as significantly correlated (Supplementary Methods). We observed a reduced FPR of 0.04 with 400 of 9900 pairs identified as significantly correlated which is within expectation based on the level of significance as p-value < 0.05 (Figure 1f).

While identifying that a gene is significantly negatively correlated across datasets can be sufficient to classify that this gene as spatially differential across the whole tissue, regions of similarity in magnitude may still be present. Likewise, genes that are significantly positively spatially correlated can be considered similar with respect to correlation but still change substantially in magnitude. Therefore, as an orthogonal visualization technique, STcompare’s spatial fold change test quantifies an overall spatial similarity (Figure 1c-d). To stabilize spatial similarity estimates, STcompare prefilters low-expression spatial locations and genes using minimal expression and coverage thresholds. It then computes the log_2_ fold change at each valid locations, annotates a location as similar if the absolute value of log_2_ fold change is within a predefined threshold, and quantifies an overall similarity score *S* for a gene as the proportion of similar locations relative to all valid locations (Supplementary Methods).

## Results

### STcompare provides insights distinct from other spatial comparison methods as demonstrated on

#### simulated data

As described previously, we simulated ST datasets where one pair of genes, A and B, had different radial expression patterns but the same mean expression, while the other pair, A and C, had same radial expression patterns but different mean expression to demonstrate that bulk DGE and SVG analysis results do not reflect changes or lack of changes in spatial expression patterns (Figure 1a). In comparison, based on both the spatial correlation and spatial fold-change quantifications from STcompare (Supplementary Methods), the gene expression patterns A and B were classified as significantly differently spatially patterned (a significant negative correlation coefficient) with 54% of locations at similar magnitude (similarity score of 0.543). Likewise, gene expression patterns A and C were classified as significantly similarly spatially patterned (a significant positive correlation coefficient) with 0% of locations at similar magnitude (similarity score of 0). In this manner, STcompare enables two differential spatial expression comparison tests that are distinct from bulk DGE and SVG analysis.

While traditional bulk DGE analysis is not spatially-aware and traditional spatially variable analysis is not informative for comparing between datasets as we have demonstrated, others have also developed methods to address these limitations. For example, STdiff identifies differentially expressed genes among tissue niches by applying non-spatial and spatial mixed models while also accounting for spatial autocorrelation to reduce false positives^16^. SPADE can identify spatially variable genes that are different between groups using Gaussian process regression machine learning models^22^. SPaSE identifies significantly pathologically remodeled regions by using optimal transport to compute a pathological score for each spot of a diseased ST dataset in comparison to a healthy ST dataset^23^. To evaluate whether these methods can address the same tasks as STcompare, we applied these methods to our simulated data. For gene expression patterns A and B, in which the gene is simulated to be spatially different but not differentially expressed, STdiff’s non-spatial test does not identify the patterns as differentially expressed so the subsequent spatial test is not evaluated and STdiff does not identify the change in spatial patterning. For gene expression patterns A and C, in which the gene is simulated to be differentially expressed yet spatially similar, STdiff successfully identifies the gene as differentially expressed using both its non-spatial mixed model (p-value = 0) and spatial mixed model differential tests (p-value = 0). However, STdiff gives no indications that gene expression patterns A and C are more spatially similar than A and B. In contrast, SPADE successfully identifies the gene expression patterns A and B as spatially variable between groups (p-value = 2.6e-58). However, SPADE also identifies gene expression patterns A and C as spatially variable between groups (p-value = 6.8e-07). As such, SPADE is not able to distinguish how gene expression patterns A and B are different from A and C. Lastly, we could not use SPaSE to identify spatially different regions between gene expression patterns A and B nor between gene expression patterns A and C, as SPaSE is designed to operate on full transcriptome ST datasets and cannot calculate a non-zero pathological score when used to compare a pair of genes (Supplementary Methods). These results thus demonstrate how these methods do not address the same tasks as STcompare.

### STcompare confirms high spatial correspondence in real ST data of biological replicates

To demonstrate STcompare on real ST data, we compared two biological replicates of coronal slices of the adult mouse brain from different animals but the same location with respect to bregma assayed by the single-cell resolution ST technology MERFISH (Figure 2a). Because these are biological replicates, we hypothesized that genes should be significantly positively correlated with high similarity, particularly for SVGs which exhibit coordinated expression variation within tissue structures that likely have correspondence across replicates.

**Figure 2.**
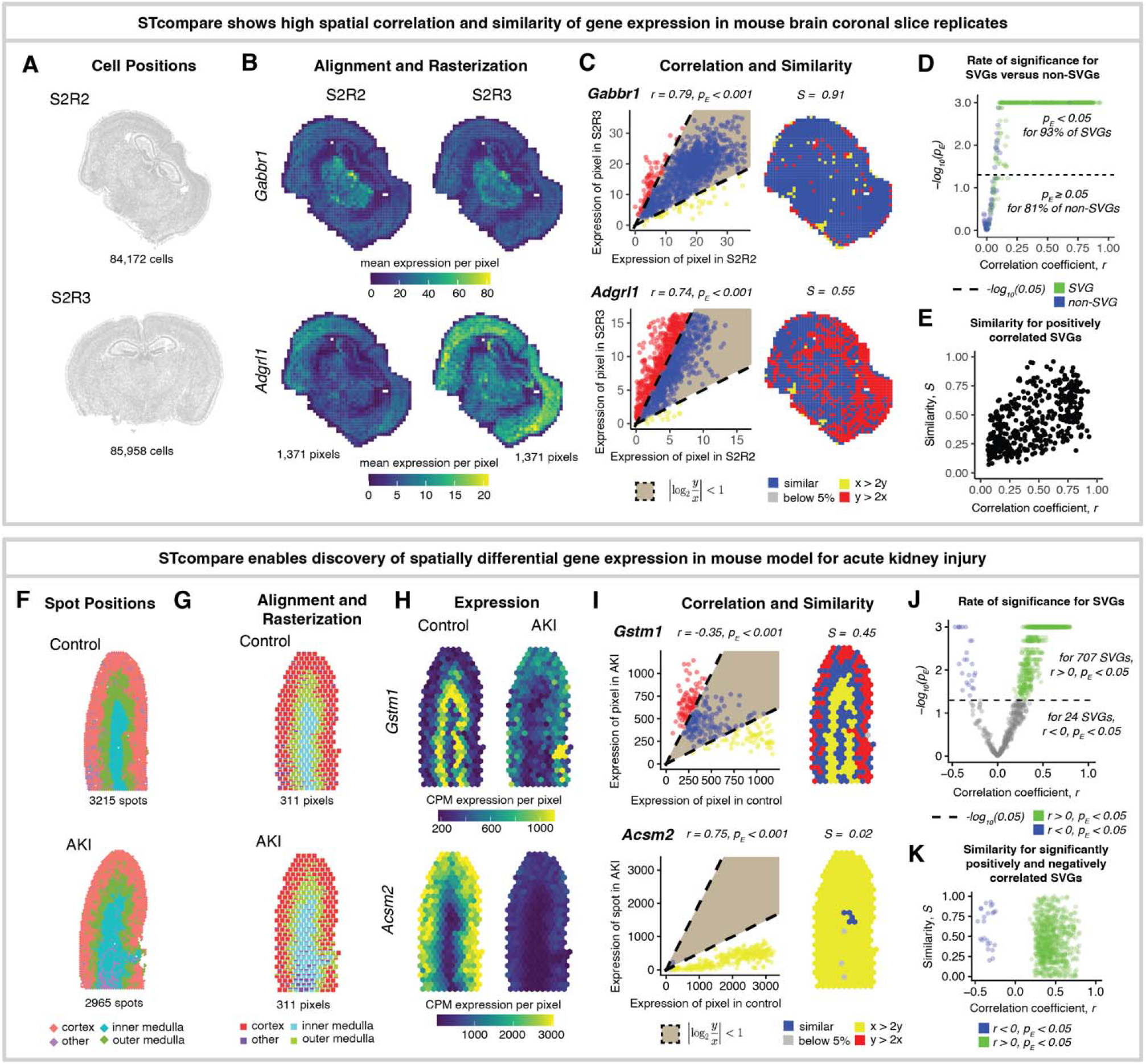
STcompare quantifies high spatial correlation and spatial similarity of gene expression patterns across biological replicates and enables discovery of spatially differential gene expression in acute kidney injury. ***A*.** Two biological replicates of coronal slices of the adult mouse brain, Slice 2 Replicate 2, S2R2, (top) and Slice 2 Replicate 3, S2R3, (bottom) assayed by single-cell resolution MERFISH *B*. Mean expression per pixel in S2R2 (left column) and S2R3 (right column) for two SVGs, *Gabbr1* (top row) and *Adgrl1* (bottom row) after alignment and rasterization. ***C***. Plots (left column) of expression in S2R2 versus S2R3 at matched pixels with correlation coefficient and empirical p-value for select similarly spatially patterned genes *Gabbr1*, r = 0.79, BH-adjusted *p*_*E*_ < 0.001 (top), and *Adgrl1*, r = 0.74, *p*_*E*_ < 0.001 (bottom). Pixels classified as either having twice as much expression in S2R3 (red), having twice as much expression in S2R2 (yellow), or similar expression (blue). Spatial visualization (right column) of spot classification with similarity scores for *Gabbr1* (S = 0.91) and *Adgrl1* (S = 0.55). Plot domains restricted to the 95^th^ quantile of pixels-based expression. **D**. For all SVGs and non-SVGs, plot of Pearson’s correlation coefficients versus the −log_10_ (*p*_*E*_) with a significance threshold of p-value > 0.05 (dashed line). 93% (384/415) of SVGs (green) identified as significantly positively correlated and 81% (55/68) of non-SVGs (blue) identified as not significantly correlated. **E**. For the 384 positively correlated SVGs, plot of the correlation coefficients versus spatial similarity score. **F**. A section from a kidney with 24-hour ischemia-reperfusion injury (bottom) and a section from a normal kidney as a control (top), assayed using the 10X Genomics Visium platform at 55um spots. Spots annotated via harmonized transcriptional clustering as spatially consistent compartments: cortex (pink), outer medulla (green), inner medulla (blue), or other (purple). **G**. Alignment and rasterization results in 311 shared pixels in control (top) and AKI (bottom). Pixels annotated with bar graphs that represent the proportion of spots from each compartment that were merged into each pixel, cortex (pink), outer medulla (green), inner medulla (blue), or other (purple). **H**. Counts per million (CPM) normalized expression in control (left column) and AKI (right column) is shown for the aligned spots for two examples, *Acsm2* (top row) and *Ndufa4* (bottom row). *I*. Plots (left column) of CPM normalized expression in control versus with AKI at matched pixels with correlation coefficient and empirical p-value for significantly negatively correlated *Gstm1*, r = −0.35, *p*_*E*_ < 0.001 (top), and significantly positively correlated *Acsm2*, r = 0.75, *p*_*E*_ < 0.001 (bottom). Pixels classified as either having twice as much expression in AKI (red), having twice as much expression in control (yellow), or similar expression (blue). The domain of plots restricted to the 95^th^ quantile of spot-based expression. Spatial visualizations (right column) of spot classifications with similarity scores for *Gstm1*, S = 0.45, (top) and *Acsm2*, S = 0.02 (bottom). *J*. For all 1046 SVGs, plot of Pearson’s correlation coefficients versus the −log_10_ (*p*_*E*_) with a significance threshold of p-value > 0.05 (dashed line). 68% (707/1046) of SVGs identified as significantly positively correlated (green) and 2% (24/1046) as significantly negatively correlated (blue). *K*. For the 707 positively correlated SVGs (green) and 24 negatively correlated SVGs (blue), plot of the correlation coefficients versus spatial similarity score.

To ensure matched spatial locations for comparison, we performed spatial structural alignment using STalign^6^ and rasterization using SEraster^13^ to obtain matching 200µm pixel locations across the two samples (Figure 2b). We identified 415 SVGs shared across both replicates by using MERINGUE^21^ to generate a list of SVGs for each replicate from the 483 genes assayed by MERFISH and then selecting the intersection of these lists. We applied STcompare’s spatial correlation test to these 415 SVGs, calculating a Pearson’s correlation coefficient *r* and a Benjamini-Hochberg (BH) adjusted empirical p-value *p*_*E*_ for each (Figure 2c, Supplementary Methods). In total, we identified 93% (384/415) of genes as significantly positively correlated and zero genes as significantly negative correlated with a BH-adjusted empirical p < 0.05 (Figure 2d), suggesting that the most SVGs have not changed in their spatial patterning, as expected. The 7% (31/415) of SVGs without significant positive correlation may reflect low expression (Supplementary Figure 1) and therefore increased susceptibility to noise (e.g., *Taar1, Ccr1, Lhcgr*), potential batch effects (e.g. *Gpr17*), or other sample-specific differences (Supplementary Figure 2).

When performing STcompare’s spatial correlation test on the remaining 68 genes not identified as SVG, 81% (55/68) were identified as not significantly positively correlated using a BH-adjusted empirical p < 0.05 (Figure 2d). The high percentage of non-SVGs identified as not significantly positively correlated is consistent with the expectation that genes not exhibiting autocorrelated spatial expression patterns are less likely to have correlated patterns across replicates. 19% (13/68) of non-SVGs were identified as significantly positively correlated across replicates. We may have observed that some non-SVGs are significantly positively correlated because we only defined genes with highly clustered patterns above a certain size as SVG and as such the non-SVG class included the remaining types of organized patterns. Three cases of non-SVGs with expression patterns that were correlated across replicates includes: (1) genes with small hotspot expression patterns (ex. *Avpr1b*), (2) genes with patterns that were organized but with high dispersion (ex. *Gpr33*), and (3) genes with low expression and increased susceptibility to noise which impacts whether the genes were classified as SVG as in both datasets (ex. *Ptafr* and *Gpr55*) (Supplementary Figure 3). Overall, this high rate of significant positive spatial correlation for SVGs and low rate for non-SVGs confirmed STcompare can quantify high spatial correspondence across biological replicates.

Of the SVGs with significant positive spatial correlation, we further used STcompare’s spatial fold change test to calculate spatial similarity with respect to change in magnitude of expression. Genes that are similarly positively spatially correlated can have different percentages of spatial similarity due to fold-change differences (Figure 2e). For example, *Gabbr1* and *Adgrl1* had similar correlation coefficients, 0.79 and 0.74, respectively, but for *Gabbr1* 91% of locations were similar whereas for *Adgrl1* only 55% of pixels were similar (Figure 2c). This lack of spatial similarity with respect to expression magnitude between these biological replicates further highlights potential batch effects and other sample-specific differences.

### STcompare enables discovery of spatially differential gene expression in acute kidney injury

Having demonstrated that STcompare works on both simulated and real ST data, we next sought to apply it to compare healthy and diseased datasets to discover potentially disease-relevant changes in spatial gene expression patterns. To this end, we focus on acute kidney injury (AKI) by comparing two previously published murine kidney sections assayed by spot-resolution 10x Visium: a section from a kidney 24 hours post initial AKI injury and a section from a sham surgery kidney as a control^24^. The two tissue sections exhibit strong structural correspondence. We highlight this by annotating the spots using harmonized transcriptional clustering, which reveals spatially consistent compartments: the inner medulla, the outer medulla, and the cortex (Figure 2f, Supplementary Figure 4). As such, this strong structural correspondence should reduce the effect of structural differences confounding discovery of disease-specific changes in spatial gene expression patterns. To further reduce the effect of confounding differences, we aligned the Visium spots by generating images from one-hot encoding the region labels and then using STalign to align the images and applying the transformation to the spots (Supplementary Methods). Next, we rasterized the aligned spots and performed counts per million (CPM) normalization (Figure 2g). Again, existing SVG analysis methods can identify genes that change from spatially variable to not or vice versa across conditions. However, genes that are classified as spatially variable in both conditions can still vary in the specific spatial pattern in each condition. For example, *Gstm1* is an SVG in both the AKI and the control datasets. But in AKI the highest expression is localized to the cortex whereas in the control the highest expression is localized the medulla (Figure 2h). To demonstrate an analysis approach that is supplementary to existing SVG analysis methods, we focus our STcompare analysis on discovering changes in spatial patterns for genes that are spatially variable in both conditions. We used MERINGUE^21^ to identify 1046 genes that are spatially variable in both datasets. We applied both STcompare’s spatial correlation test and spatial fold change test to the 1046 SVGs, calculating a Pearson’s correlation coefficient *r*, an empirical p-value *p*_*E*_, and a spatial similarity score *S* for each gene (Figure 2i). After applying BH procedure to obtain adjusted p-values, we identified 2% (24/1046) of genes as significantly differently spatially patterned (*r* < 0, *p*_*E*_ < 0.05) between AKI and control (Figure 2j, Supplementary Table 1). Additionally, we identified 68% (707/1046) of SVGs as significantly similarly spatially patterned (r > 0, *p*_*E*_ < 0.05) between AKI and control (Figure 2j, Supplementary Table 2). We further confirm that STcompare’s spatial fold-change test can provide orthogonal insights from the spatial correlation test (Figure 2k) based on the general lack of correlation between these metrics. Given such full-transcriptome spatial transcriptomics data, we also sought to evaluate and compare the results of STcompare with that of SPaSE^23^ (Supplementary Methods). Applying SPaSE identified a singular significantly pathologically remodeled region in the AKI tissue that encompassed all the spots assayed by Visium (Supplementary Figure 8), highlighting the injury affects the entire tissue but not distinguishing between cortical and medullary effects. In addition, SPaSE did not identify spatial changes for specific genes, again demonstrating how these other methods do not address the same tasks as STcompare.

Of the genes that STcompare identified as spatially differentially expressed, we used visualizations of the gene expression and the similarity scores to observe that some genes changed their spatial expression patterns in a compartment-specific manner, with expression decreasing in one compartment but increasing in a different compartment. *Gstm1, Cbr1, Psap, Acadl*, and *Nop10*, genes involved in metabolic pathways, which are not specific to a singular cell-type, decreased expression in medulla and increased expression in the cortex in AKI compared to control (Figure 2i, Supplementary Figure 5). We speculate this spatial pattern change may be indicative of metabolic reprogramming as expression of *Gstm1 and Cbr1* may suggest generation of the anti-oxidant glutathione to confer protection against AKI mediated oxidative stress^25^, *Psap* and *Acadl* are essential for lipid breakdown^26,27^, and *Nop10* codes for a nucleolar protein involved in metabolic stability of RNA^28^. Further, genes involved in oxidative phosphorylation (OXPHOS), such as *Ech1, Ndufs6, Mpc, Atp5l, Cox7b, and Cox8a*, have decreased expression in the medulla but comparable expression in the cortex in AKI relative to the control (Supplementary Figure 6, Supplementary Table 1). This sustained expression of OXPHOS related genes within the cortex of AKI tissue might be indicative of an injured tissue state slowly returning to normalcy at 24 hours post initial AKI injury with the blood flow allowed to go back into the kidney tissue. Other genes that we observed with a decreased expression in the medulla but comparable expression in the cortex in AKI relative to the control include *Vamp8*, a vesicle-associated membrane protein critical for lysosomal fusion, *Nceh1*, a cholesterol ester hydrolase important for lipid metabolism, and *Csad*, a cysteine sulfinic acid decarboxylase essential for taurine biosynthesis (Supplementary Figure 7). These different changes suggests that injury response in this mouse model of AKI is spatially organized, occurring in a tissue compartment-specific manner perhaps due to compartment-specific metabolic demands and preferences.

The genes that STcompare identified as significantly positively correlated that also have relatively higher similarity scores include ribosomal proteins *Rpl35a* (*r* = 0.45, *p*_*E*_ < 0.001, *S* = 0.97) and *Rpl41* (*r* = 0.50, *p*_*E*_ < 0.001, *S* = 0.96), suggesting that these genes, which are ubiquitous and essential for protein synthesis, may not be impacted by AKI at this evaluated severity or timeframe in the disease process (Supplementary Figure 9). In contrast, some of the significantly positively correlated genes did not change in spatial patterning but did change in magnitude. For example, *Acsm2* (*r* = 0.75, *p*_*E*_ < 0.001, *S* = 0.02), a gene encoding an enzyme in the fatty acid oxidation pathway, and *Slc34a1* (*r* = 0.75, *p*_E_ < 0.001, *S* = 0.01), a gene in the solute carrier family, are normally specifically expressed in the proximal tubule cells within the cortex^29^. In AKI compared to control, these cell-type specific genes remained localized to the cortex, but expression was greatly reduced (Figure 2i, Supplementary Figure 9). This is consistent with proximal tubule-specific consequence in AKI^30^. Further, genes that are localized to the cortex and protective against oxidative stress and cell death (apoptosis and ferroptosis) such as *Nox4*^*31*^ (*r* = 0.53, p_*E*_ < 0.001, *S* = 0.31)and *Gpx1*^*32*^ (*r* = 0.79, p_*E*_ < 0.001, S = 0.30) also decreased in AKI (Supplementary Figure 10), while the genes involved in the fibrotic process such as *Spp1*^*33*^ (*r* = 0.55, p_*E*_ < 0.001, *S* = 0) and *Tmsb10*^*34*^ (r 0.49, *p*_*E*_ 0.004, *S* 0.06) ^*35*^increased in cortex in AKI (Supplementary Figure 11). These findings suggest that damaged proximal tubule cells may remain spatially in their cortical niche but transcriptionally shift to prioritize fibrotic processes over normal function. Thus, application of STcompare to identify spatially differential genes in AKI identifies molecular processes spatially dysregulated in a disease model, improving characterization of disease processes.

## Discussion

Comparative spatial transcriptomics data analysis is needed to characterize differentially spatially patterned genes for structurally matched tissues across axes of interest such as health versus disease. Here, we presented STcompare, which enables two differential spatial expression comparison tests: a spatial correlation test and a spatial fold change test. The spatial correlation test identifies gene expression patterns that have shifted in spatial dependencies across datasets by defining differently spatially patterned genes or similarly spatially patterned genes as those with statistically significant negative or positive Pearson’s correlation coefficients, respectively, while controlling for spatial autocorrelation that may inflate spatial statistics. Since correlation alone may miss differences in the absolute magnitude of spatial expression, the spatial fold change test measures similarity with respect to magnitude. By calculating the fold-change at each individual spatial location, we are also able to visualize regions of low and high similarity to identify region-specific changes in magnitude.

We evaluate the performance of STcompare using simulated data as well as real ST data of biological replicates to demonstrate that STcompare enables insights distinct from bulk DGE analysis and recapitulate the expected high spatial correspondence of gene expression across replicates. Furthermore, we apply STcompare to discover potentially disease-relevant changes in spatial gene expression patterns by comparing ST data from mouse kidney with acute kidney injury versus a control mouse kidney.

Beyond the applications we have demonstrated, STcompare can also be applied to evaluate technological differences when applied to structurally matched tissues assayed by different ST technologies. Additionally, STcompare can be applied to evaluate cell-type correspondence by comparing cell-type proportions at matched locations^6^. Furthermore, STcompare can be used to compare across omics modalities such as gene and protein expression when applied to serial tissue sections assayed by different spatial omics technologies.

While we propose additional applications of STcompare, we acknowledge that there are limitations to the applications of STcompare. Both the spatial correlation test and spatial fold-change test of STcompare are dependent on the datasets under comparison having inherent structural similarity. In the cases in which datasets need to be aligned, STcompare is limited by how well the datasets can be aligned, and, if necessary, rasterized, to maximize the structural correspondence and create comparable spatial locations. To assess the sensitivity of results to alignment errors, we apply STcompare to evaluate the MERFISH biological replicates with less precise affine alignments compared to STalign alignment. We find STcompare’s correlation test (particularly with respect to the significance testing) and the fold-change similarity test are robust to slight change in alignments (Supplementary Figure 12). However, in general, when structural alignment is poor, the interpretation of STcompare results may become challenging, as any statistically significant differences could reflect underlying alignment error.

Extending beyond the analyses presented here, while we classified only genes with statistically significant negative Pearson’s correlation coefficients as differently spatially patterned genes, representing a conservative approach, a less conservative approach would have been to define any gene that does not have a statistically significant positive correlation as differently spatially patterned, including genes with nonsignificant small magnitude Pearson’s correlation coefficients. Users of STcompare are provided all correlation coefficients and corresponding p-values without interpretation so they can adjust definitions as fit for their biological questions of interest. Additionally, the precision and accuracy of the empirical p-values calculated in this test are limited by the assumptions and methods used when generating the permutations. Firstly, while increasing the number of permutations could increase empirical p-value precision, doing such would also increases computational time. To increase computationally efficiency, we implement parallelization across permutations, and we provide a function for users to filter out genes for which the analytical or empirical p-value is much greater than 0.05 before increasing the number of permutations, as increasing precision would not affect interpretation of significance of those cases. In general, runtime for the spatial correlation test scales proportionally with the number of genes and pixels analyzed (Supplementary Figure 13, Supplementary Methods). Secondly, by employing gaussian kernel smoothing to generate permutations we assume that autocorrelated patterns in gene expression can be abstracted to a radial shape. If this assumption fails for a gene, the variograms of the permutations generated from Gaussian kernels will not fit well to the variogram of the gene and users may observe high variance for 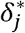 for *j* [1, ‖, *B*], which is the optimal *δ* ^*^ (percentage of neighbors that should be within the area enclosed by the radius of the Gaussian kernel smoothing) for each of the B permutations of a gene. If users observe high variance for 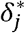, they should modify *δ* by increasing the range of the numbers provided. If 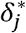 continues to have high variance with an expanded range for *δ*, then it is possible that the assumption that the autocorrelated pattern can be mimicked with a Gaussian kernel fails for that gene. To investigate further, the user can plot a variogram of the gene of interest and observe if the relationship between distance of points and difference in gene expression appears Gaussian. If the Gaussian assumption is not appropriate, STcompare could be modified in the future to accommodate other kernels or geometries for generating permutations for genes with layered or more complex patterns. Lastly, our ability to generate permutations that have similar degrees of autocorrelation as the original data is limited by the metric which we use to evaluate the difference in degree of autocorrelation between datasets. We employ variograms, which plot a measure of difference between spatial locations as a function of the distance between them, as a means of measuring autocorrelation but variograms are less precise at greater distances because as the distance increases there are fewer pairs of spatial locations with that distance between them. Since the goal is to minimize the difference between the variogram of X and those of its permutations, we choose to subset to the 25th percentile of the variograms by default to improve robustness. Users of STcompare could try increasing the percentile to get better permutations but they would need to identify a novel means for determining if the resulting permutations are a better match to the original data as increasing the percentile will inherently increase the sum of squared errors between the variograms subset to that percentile. Regarding the spatial similarity test, the three parameters that can be user defined are the fold change threshold, the minimum gene expression threshold, and minimum pixels threshold. Since a spatial location is classified as similar if the absolute value of log_2_ fold change is below a predefined threshold *b*, users can make this classification more or less conservative by making this threshold lesser or greater, respectively. Since inclusion of pixels with low expression or genes with too many low expression pixels may result in inflated and variable similarity scores, respectively, users can define a minimum gene expression threshold and a minimum pixels threshold. The spatial similarity test is also limited in that we do not test for significance. However, we intend for the similarity scores to primarily be used as an exploratory visualization aid to show regions of similar expression magnitude based on fold-change. In this manner, the similarity score is complementary to non-spatial differential expression methods that have tests for significance.

We anticipate that STcompare may be integrated with other complementary spatial analysis methods. As we have shown, STcompare can supplement SVG analysis methods that identify changes in SVG status^22^ by further detecting pattern shifts when a gene remains spatially variable across conditions.

Likewise, STcompare may be applied in conjunction with spatial domains detection methods^12,36^. In particular, since the correlation test of STcompare is subject to cases of Simpson’s paradox, in which negative correlations that are statistically significant for individual domains can lose significance or become positive correlations when the domains are aggregated, restricting STcompare analysis to specific spatial domains can, in some cases, clarify correlation results. However, we would not advise always limiting analysis to individual spatial domains as many spatial processes may span across spatial domain. Accordingly, STcompare also complements transcriptional clustering analyses that define spatially patterned cell-types and perform cell-type-specific differential expression^16,37^ by capturing processes that span across multiple cell-types. Overall, STcompare expands the suite of spatial-omics analysis methods, providing complementary tests that can be combined or used in parallel with existing methods to enable more thorough comparative analyses of spatially resolved -omics datasets and reveal differential spatial molecular patterns across various physiological and technological axes.

## Availability of data and materials

The MERFISH mouse brain datasets analyzed during the current study are available in a Zenodo repository at https://zenodo.org/records/10724029. The Visium mouse kidney datasets analyzed during the current study are available in a Zenodo repository at https://zenodo.org/records/17676992. Data generated during this study are included in this article and its supplementary information files.

## Acknowledgements

This material is based upon work supported by the National Science Foundation under CAREER-2047611 (J.F., K.C., V.J.) and the HuBMAP Integration, Visualization, and Engagement (HIVE) Initiative under Award Number OT2-OD033760 (J.F., S.S., R.d.S.P.). We thank YooJin Chloe Sheen for her exploratory analysis of *δ* values. We thank Robert Tibshirani and Trevor Hastie for earlier discussions on spatial statistics that motivated this work.

## Author contributions

J.F. and K.C. conceptualized the study. K.C. and V.J. developed the computational software and performed bioinformatics analyses. R.d.S.P. beta-tested the software. S.S., R.M, and H.R. contributed biological expertise to the interpretation of the kidney results. K.C., V.J., and J.F. wrote the manuscript. All authors reviewed the results and approved the final version of the manuscript.

## REFERENCE

1. Bressan, D., Battistoni, G. & Hannon, G. J. The dawn of spatial omics. Science (1979). 381, (2023).

2. Moffitt, J. R., Lundberg, E. & Heyn, H. The emerging landscape of spatial profiling technologies. Nat. Rev. Genet. 23, 741–759 (2022).

3. Gulati, G. S., D’Silva, J. P., Liu, Y., Wang, L. & Newman, A. M. Profiling cell identity and tissue architecture with single-cell and spatial transcriptomics. Nat. Rev. Mol. Cell Biol. 26, 11–31 (2025).

4. Jain, S. & Eadon, M. T. Spatial transcriptomics in health and disease. Nature Reviews Nephrology | 20, 659–671 (2024).

5. Bonev, B. et al. Opportunities and challenges of single-cell and spatially resolved genomics methods for neuroscience discovery. Nat. Neurosci. 27, 2292–2309 (2024).

6. Clifton, K. et al. STalign: Alignment of spatial transcriptomics data using diffeomorphic metric mapping. Nat. Commun. 14, 8123 (2023).

7. Zeira, R., Land, M., Strzalkowski, A. & Raphael, B. J. Alignment and integration of spatial transcriptomics data. Nat. Methods 19, 567–575 (2022).

8. Tang, Z. et al. Search and match across spatial omics samples at single-cell resolution. Nat. Methods 21, 1818–1829 (2024).

9. Jones, A., Townes, F. W., Li, D. & Engelhardt, B. E. Alignment of spatial genomics data using deep Gaussian processes. Nat. Methods 20, 1379–1387 (2023).

10. Bergenstråhle, J., Larsson, L. & Lundeberg, J. Seamless integration of image and molecular analysis for spatial transcriptomics workflows. BMC Genomics 21, 1–7 (2020).

11. Li, H. et al. SANTO: a coarse-to-fine alignment and stitching method for spatial omics. Nat. Commun. 15, (2024).

12. Zhou, X., Dong, K. & Zhang, S. Integrating spatial transcriptomics data across different conditions, technologies and developmental stages. Nat. Comput. Sci. 3, 894–906 (2023).

13. Aihara, G. et al. SEraster: a rasterization preprocessing framework for scalable spatial omics data analysis. Bioinformatics 40, (2024).

14. Viladomat, J., Mazumder, R., Mcinturff, A., Mccauley, D. J. & Hastie, T. Assessing the Significance of Global and Local Correlations under Spatial Autocorrelation: A Nonparametric Approach. Biometrics 70, 409–418 (2014).

15. Zhu, J., Sun, S. & Zhou, X. SPARK-X: non-parametric modeling enables scalable and robust detection of spatial expression patterns for large spatial transcriptomic studies. Genome Biol. 22, (2021).

16. Ospina, O. E. et al. Differential gene expression analysis of spatial transcriptomic experiments using spatial mixed models. Sci. Rep. 14, 10967 (2024).

17. Edsgärd, D., Johnsson, P. & Sandberg, R. Identification of spatial expression trends in single-cell gene expression data. Nat. Methods 15, 339–342 (2018).

18. Weber, L. M., Saha, A., Datta, A., Hansen, K. D. & Hicks, S. C. nnSVG for the scalable identification of spatially variable genes using nearest-neighbor Gaussian processes. Nat. Commun. 14, 4059 (2023).

19. Hu, J. et al. SpaGCN: Integrating gene expression, spatial location and histology to identify spatial domains and spatially variable genes by graph convolutional network. Nat. Methods 18, 1342–1351 (2021).

20. Svensson, V., Teichmann, S. A. & Stegle, O. SpatialDE: identification of spatially variable genes. Nat. Methods 15, 343–346 (2018).

21. Miller, B. F., Bambah-Mukku, D., Dulac, C., Zhuang, X. & Fan, J. Characterizing spatial gene expression heterogeneity in spatially resolved single-cell transcriptomic data with nonuniform cellular densities. Genome Res. 31, 1843–1855 (2021).

22. Qin, F. et al. Spatial pattern and differential expression analysis with spatial transcriptomic data. Nucleic Acids Res. 52, e101–e101 (2024).

23. Rahman, M. N. et al. SPaSE: Spatially resolved pathology scores using optimal transport on spatial transcriptomics data. Cell Syst. 16, 101301 (2025).

24. Gharaie, S. et al. Single cell and spatial transcriptomics analysis of kidney double negative T lymphocytes in normal and ischemic mouse kidneys. Sci. Rep. 13, 20888 (2023).

25. van der Pol, A. et al. Accumulation of 5-oxoproline in myocardial dysfunction and the protective effects of OPLAH. Sci. Transl. Med. 9, (2017).

26. Meyer, R. C., Giddens, M. M., Coleman, B. M. & Hall, R. A. The protective role of prosaposin and its receptors in the nervous system. Brain Res. 1585, 1–12 (2014).

27. Chiba, T. et al. Loss of long-chain acyl-CoA dehydrogenase protects against acute kidney injury. https://doi.org/10.1172/jci (2025) doi:10.1172/jci.

28. Elsharawy, K. A. et al. Nucleolar protein 10 (NOP10) predicts poor prognosis in invasive breast cancer. Breast Cancer Res. Treat. 185, 615–627 (2021).

29. Gisler, S. M. et al. PDZK1: I. A major scaffolder in brush borders of proximal tubular cells11See Editorial by Moe, p. 1916. Kidney Int. 64, 1733–1745 (2003).

30. Shan, D. et al. Dynamic cellular changes in acute kidney injury caused by different ischemia time. iScience 26, 106646 (2023).

31. Li, F. et al. Single-cell analysis of proximal tubular cells with different DNA content reveals functional heterogeneity in the acute kidney injury to chronic kidney disease transition. Kidney Int. 108, 90–104 (2025).

32. Zhang, Y., Handy, D. E. & Loscalzo, J. Adenosine-Dependent Induction of Glutathione Peroxidase 1 in Human Primary Endothelial Cells and Protection Against Oxidative Stress. Circ. Res. 96, 831–837 (2005).

33. Ding, H. et al. Kidney fibrosis molecular mechanisms Spp1 influences fibroblast activity through transforming growth factor beta smad signaling. iScience 27, 109839 (2024).

34. Qin, Z. et al. Single-Cell Sequencing Uncovers a TMSB10-Expressing Fibroblast Subpopulation Driving Renal Fibrosis in Diabetic Nephropathy. Diabetes, Metabolic Syndrome and Obesity Volume 18, 4913–4929 (2025).

35. Kobayashi, H. et al. Neuroblastoma suppressor of tumorigenicity 1 is a circulating protein associated with progression to end-stage kidney disease in diabetes. Sci. Transl. Med. 14, (2022).

36. Kang, L., Zhang, Q., Qian, F., Liang, J. & Wu, X. Benchmarking computational methods for detecting spatial domains and domain-specific spatially variable genes from spatial transcriptomics data. Nucleic Acids Res. 53, 303 (2025).

37. Cai, P., Robinson, M. D. & Tiberi, S. DESpace: spatially variable gene detection via differential expression testing of spatial clusters. Bioinformatics 40, (2024).

