## Supplementary Material for "STcompare: comparative spatial transcriptomics data analysis of structurally matched tissues to characterize differentially spatially patterned genes"

**Supplementary Methods**

1. STcompare overview

To identify pairwise spatially differentially expressed genes using STcompare, we assumed datasets have been structurally aligned to resolve $N$ matched spatial locations.

*Overview of spatial correlation method*

To identify gene expression patterns that have shifted in spatial dependencies across datasets, we define a gene as differently spatially patterned or similarly spatially patterned if it has a statistically significant negative or positive Pearson’s correlation coefficient, $r$ , respectively. Given $x$ is the expression of a gene in the first dataset and $y$ is the expression of a gene in the second dataset, the equation for Pearson’s correlation coefficient is (1)

$$r=\frac{\sum\left( x_{i}-\bar{x} \right)\left( y_{i}-\bar{y} \right)}{\sqrt{\sum\left( x_{i}-\bar{x} \right)^{2}\sum\left( y_{i}-\bar{y} \right)^{2}}}$$

where $i = \left[ 1, \ldots N \right]$ matched spatial locations, $x_{i}$ is the expression in the first dataset at the $i$*-*th spatial location, $\bar{x}$ is the mean expression in the first dataset across all spatial points, and likewise, $y_{i}$ is the expression in the second dataset at the $i$*-*th spatial location, and $\bar{y}$ is the mean expression in the second dataset across all spatial locations. To generate an analytical p-value, the Pearson’s correlation coefficient is evaluated with a two-sided alternative hypothesis and with a test statistic following a t- distribution with $N-2$ degrees of freedom assuming the values in the data are independently and normally distributed, as implemented in the default settings of the `cor.test()` function of the R package *stats* version 4.5.0^1^. Since spatially patterned genes often have spatial autocorrelation that violates the assumption of the analytical statistical tests that values in the data are independently distributed, we also calculate an empirical p-value by generating an empirical null distribution to evaluate the Pearson’s correlation coefficient against. The approach for generating this empirical p-value was previously described in Viladomat et al^2^ and the specifics of this implementation are described as follows.

Given expression of a gene in two datasets represented as $X$ and $Y$, we create permutations of $X$. These permutations are intended to have the same degree of autocorrelation as the original $X$. The degree of autocorrelation of $X$ is determined by calculating an empirical variogram, which gives a measure of difference between points in $X$ as a function of the Euclidean distance between points in $X$. Calculation of variograms was implemented using `variog` function from R package *geoR* version 1.9.5^3^ with `estimator.type = "classical"` so that the empirical variogram value would be estimated using classical method of moments estimator and `option = "bin"` so that binned data would be returned as output. Given that a variogram gets noisier as distances between points increase, we set `max.dist`, which is the upper limit of the domain of the variogram, to the 25th percentile of the distances between points. Defaults were used for all other arguments of `variog`. We refer to the variogram of $X$ as the “target variogram”.

We define permutations of $X$ as $X_{j}$ for $j = [1, ..., B]$, where $B$ is the number of permutations (default $B$ = 100). Initially, a permutation $X_{j}$ is generated by randomly shuffling $X$ via taking a sample of size $N$, i.e the length of $X,$ from the elements of $X$ without replacement. To make $X_{j}$ have autocorrelated values, we applied Gaussian kernel smoothing to the shuffled $X_{j}$. This smoothing was implemented using the `locfit` and `lp` functions from R package *locfit* version 1.5.9.12^4^ which performs local weighter regression. For the `locfit` function we set `kern = “gauss”` to set a Gaussian as the weight function and increased the `maxk` to 300. Defaults were used for all other arguments of `locfit`. We used the `lp` function to model $X_{j}$ as a function of the spatial coordinates using a zero-degree polynomial. The `nn` argument of `lp`, the nearest neighbor component, determines the radius of the Gaussian kernel smoothing. The value set to `nn` is the percentage of neighbors that should be within the area enclosed by the radius of the gaussian kernel smoothing. We define this value as $\delta.$ Instead of choosing one numeric for $\delta$, we allow this parameter to be optimized over a range since spatial patterns can have degrees of autocorrelation at different scales (default $\delta= [0.1, 0.2, 0.3, 0.4, 0.5, 0.6, 0.7, 0.8, 0.9])$. Given that $X_{j}$ is a function of $\delta$, to get each $X_{j}$ for $j = [1, ..., B]$ we choose the $\delta^{*}$ that minimizes the sum of squared errors between the variograms of $X$ and $X_{j}$ such that there is $\delta_{j}^{*}$ for $j = [1, ..., B]$. Users may observe high variance for $\delta_{j}^{*}$ if the true optimal $\delta^{*}$ is not within the range of values inputted as $\delta$. If users observe high variance for $\delta_{j}^{*}$, they should modify $\delta$ by increasing the range of the numbers provided. To be computationally efficient, we advise users to modify $\delta$ only for the genes for which there is high variance for $\delta_{j}^{*}$ for $j = [1, ..., B]$, instead of increasing number of values provided as $\delta$ for all genes.

Next, computing the Pearson’s correlation coefficient, $r_{j}$, for $Y$ and each $X_{j}$ for $j = [1, ..., B]$ results in an empirical null distribution. The equation for this Pearson’s correlation coefficient is (2)

$$r_{j}= \frac{\sum_{i=1}^{N} (X_{i,j}-\bar{X_{j}})(Y_{i}-\bar{Y})}{\sqrt{\sum_{i=1}^{N} {(X_{i,j}-\bar{X_{j}})}^{2}\sum_{i=1}^{N} {(Y_{i}-\bar{Y})}^{2}}}$$

where $i = \left[ 1, \ldots N \right]$ spatial points, $j = \left[ 1, \ldots B \right]$ permutations, $X_{i, j}$ is the expression at the $i$*-*th spatial point in the $j$*-*th permutation, $\bar{X_{j}}$ is the mean expression across all spatial points in $j$*-*th permutation, $Y_{i}$ is the expression in the second dataset at the $i$*-*th spatial point, and $\bar{Y}$ is the mean expression in the second dataset across all spatial points. Finally, a two tailed empirical p-value, $p_{E}$, is calculated as (3) and (4)

$g\left( r_{j}, r \right)= \left\{ \begin{aligned} 1, \left| r_{j} \right|>\left| r \right| \\ 0, \text{otherwise} \end{aligned} \right.$

$$p_{E}= \frac{1}{B}\sum_{j=1}^{B} g\left( r_{j}, r \right)$$

i.e., the proportion $r_{j}$ that has an absolute value greater than the absolute value of $r$, where $r_{j}$ is the correlation coefficients for $Y$ and each $X_{j}$ and $r$ is the correlation coefficient for $X$ and $Y$ .

We calculate $p_{E}$when $x$ corresponds to first dataset and $y$ the second dataset such that the first dataset is permuted and then we calculate $p_{E}$again after swapping the datasets so that $x$ is the second dataset and $y$ is the first dataset such that the second dataset is the permuted one. We report $p_{E}$ as the greater (ie. more conservative) value from the two scenarios.

*Overview of spatial similarity method*

To identify regions in gene expression patterns that change substantially in magnitude across datasets, which can occur even if the spatial patterns are positively correlated, we define a matched spatial location as similar if the absolute log_2_ fold change of expression at that spatial location across the two datasets is less than a threshold $b$ (5)

$$f\left( x_{i},y_{i} \right)=\left\{ \begin{aligned} 1, &\left| \log_{2} \frac{y_{i}}{x_{i}} \right|<b \\ 0, &\text{otherwise} \end{aligned} \right.$$

where $x_{i}$ is the expression in the first dataset at the $i$*-*th spatial location, $y_{i}$ is the expression of in the second dataset at the $i$*-*th spatial location, and $b$ is the threshold (default = 1). To capture the overall spatial similarity, we further calculate a similarity score, $S$ as the proportion of spatial locations that are similar compared to the total number of spatial locations, given by equation (6)

$$S= \frac{1}{N_{q}}\sum_{i=1}^{N_{q}} f(x_{i},y_{i})$$

where $x_{i}$ is the expression in the first dataset at the $i$*-*th spatial location, $y_{i}$ is the expression in the second dataset at the $i$*-*th spatial location, and $N_{q}$ is the total number of spatial locations for which either $x_{i}$ is greater than $q$ of $x$, or $y_{i}$ is greater than $q$ of $y$ where $q=0.05$ by default (i.e. 5% or the 5^th^ quantile). This selection criterium for $N_{q}$ is employed before calculating the similarity score to remove spatial locations with low expression across both datasets.

A similarity score is not calculated and `spatialSimilarity` returns `NA` for lowly rarely expressed genes, specifically those genes for which $N_{q}$ is less than a minimum spatial locations threshold (default 10% of the total spatial locations). The similarity score, $S$, can be interpreted as the fraction of the spatial locations that are similar.

To visualize the spatial locations classified as similar we use the aforementioned equation (5) but assign a color. Additionally, to visualize the sign, i.e. direction, of the log2 fold-change for spatial locations which are not similar, we assign two additional different colors using the following paradigm (7)

$$f\left( x_{i},y_{i} \right)=\left\{ \begin{aligned} \text{blue}, &\left| \log_{2} \frac{y_{i}}{x_{i}} \right|<b \\ \text{yellow}, &\log_{2} \frac{y_{i}}{x_{i}}<-b \\ \text{red}, &\log_{2} \frac{y_{i}}{x_{i}}>b \end{aligned} \right.$$

For thoroughness, the percentage of spatial locations in each of these two dissimilar classes is also returned in our code implementation of this method.

2. Simulated Data with Positively and Negatively Correlated Spatial Patterns

*Data Generation for Simulated Data with Positively and Negatively Correlated Spatial Patterns*

To simulate pairs of positively correlated and negatively correlated genes, we generated three kidney shaped datasets, A, B, and C, each with one spatially patterned gene. Each simulated datasets began with simulating cells as 5000 random points with $(x,y)$ coordinates generated within a bounding box from $x \epsilon[-3,3]$ and $y \epsilon[-3,3]$. Then, to select only cells within the shape of a kidney, a cell at $(x,y)$ was only kept if its Euclidean distance from the origin was less than or equal a distance specified by the following equations:

$$\sqrt{x^{2}+ y^{2}} \leq1-2\sin\left( \theta\right)+\cos\left( \theta^{2} \right)$$

$$\theta=\text{atan2}(y,x)$$

Cells were then flipped over the $y=x$ axis such that $\left( x,y \right)=(y+0.75, x)$. We distributed gene expression values for each dataset, $G_{A},G_{B}, \text{and} G_{C}$, with the following equations

$$G_{A}=\left( 4x \right)^{2}+\left( 2y \right)^{2}+Z_{A}+10$$

$$G_{B}=10-\left( \left( 4x \right)^{2}+\left( 2y \right)^{2}+Z_{B} \right)+63$$

$$G_{C}={2\left( 4x \right)}^{2}+{2\left( 2y \right)}^{2}+{2Z}_{C}+20$$

Where $Z$is a list of random deviates from a normal distribution with mean of 20 and standard deviation of $\sqrt{10}$generated using the `rnorm` function from R package *stats* version 4.5.0. The length of $Z$for each dataset is the number of cells within the kidney shape for that dataset. Each of the gene expression vectors $G_{A},G_{B}, \text{and} G_{C}$ were then transformed by subtracting the minimum of the aggregate of all datasets, i.e. ${\text{min}(G}_{A},G_{B}, G_{C})$, so that the new minimum was zero.

*Bulk Differential Expression Analysis of Simulated Data with Correlated Spatial Patterns*

Log_2_ fold-change was calculated as the mean of the expression of the gene in one dataset divided by the mean of the expression of the gene in another dataset and then log_2_ transformed. Two-sided two-sample Wilcoxon rank sum tests were performed to test if the distributions differ by a location shift greater than 0 with p-values computed through normal approximation as implemented in `wilcox.test` function of R package *stats* version 4.5.0^1^.

*Spatially Variable Gene Analysis of Simulated Data with Correlated Patterns*

We performed spatially variable gene (SVG) analysis on the simulated datasets A, B, and C using Moran’s I as implemented in R package *MERINGUE* version 1.0^5^ with the default parameters.

*Rasterization*

For the three correlated simulated datasets, we formatted the position and gene expression from each dataset into `SpatialExperiment` objects using the R package *SpatialExperiment* version 1.18.1^6^. These three `SpatialExperiment` objects were rasterized into hexagonal pixels with resolution = 0.2 using R package *SEraster* version 1.0.0^7^, such that each pixel value is the mean of gene expression of all simulated cells within the boundaries of the pixel. Rasterizing the three spatial experiments together as a list ensured that pixel grids (ie. spatial locations) of each datasets have the same boundaries and pixels at the same coordinates across datasets have the same names across datasets.

*Calculation of Spatial Correlation*

For the negatively correlated pair, simulated datasets A and B, and the positively correlated pair, simulated datasets A and C, we used the `spatialCorrelationGeneExp` function to calculate the correlation coefficient (and the associated empirical p-value) to compare the rasterized gene expression in the pair. We used the default parameters in which $B=100$ permutations and $\delta= [0.1, 0.2, 0.3, 0.4, 0.5, 0.6, 0.7, 0.8, 0.9]$.

*Calculation of Spatial Similarity*

We used the `spatialSimilarity` function to calculate the similarity score to compare the rasterized gene expression within each pair of simulated correlated datasets, A/B and A/C. Since the simulated data does not have as many pixels with zero expression as real data, we did not use the default $q=0.05$ to remove the pixels below the 5^th^ quantile of either dataset. Instead, we set `t1 = 0` and `t2 = 0`, to use all pixels with non-zero expression in the either first or second datasets to calculate the similarity score, effectively letting $N_{q} =N$ . All other parameters were left as default.

3. Simulated Data with No Correlated Spatial Patterns

*Data Generation for Simulated Data with No Correlated Spatial Patterns*

To have simulated genes with no correlation, we generated 100 kidney shaped datasets, each with one independently randomly generated spatially patterned gene. Each simulated datasets began with simulating cells as $N=5000$ random points with $(x,y)$ coordinates generated within a bounding box from $x \epsilon[-3,3]$ and $y \epsilon[-3,3]$. The gene expression value $G_{i}$ for the $i$*-*th simulated cell was generated from realizations of Gaussian random fields with independent and identically distributed errors. Specifically, a Matern covariance matrix, $\text{Cov}_{ij}$, of size $N$ x $N$ was computed using the following equation

$$\text{Cov}_{ij}=\frac{2^{1- \nu}}{\Gamma(\nu)}*\left( \frac{h\sqrt{2\nu}}{\kappa} \right)^{\nu}* K_{\nu}\left( \frac{h\sqrt{2\nu}}{\kappa} \right)$$

$$h= \sqrt{{(x_{i}-x_{j})}^{2}+ {(y_{i}-y_{j})}^{2}}$$

where $(x_{i}, y_{i})$ is the location of the $i$*-*th cell for $i = \left[ 1, \ldots N \right]$, $(x_{j}, y_{j})$ is the location of the $j$*-*th cell for $j = \left[ 1, \ldots N \right]$, the smoothness parameter $\nu=0.5$, the range parameter $\kappa=0.1$, $\Gamma$ is the gamma function, and $K_{\nu}$ is the Bessel K function of order $\nu$. In the case that $i=j$, we set $\text{Cov}_{ij}=1$. The covariance matrix $\text{Cov}_{ij}$ was used to generate a multivariate normal distribution with $\mu=0$ which was sampled to produce an expression vector $W_{i}$ of length $N$, via the function `mvrnorm` from the R package *MASS* version 7.3.65^8^. To get the vector $Z_{i}$ of independently and identically distributed errors, we generated $N$ random deviates from a normal distribution with mean of 0 and standard deviation of $\sqrt{0.3}$ using the `rnorm` function from R package *stats* version 4.5.0^1^. Finally, $G_{i}= W_{i}+ Z_{i}+10$ resulted in a non-negative random expression value for $i$*-*th simulated cell.

Then, to select only cells within the shape of a kidney, cell positions were first transformed to $\left( x,y \right)=(6x-3, 6x-3)$ to take advantage of the formula already described above for shaping the other simulated dataset into a kidney. Again, a cell at $(x,y)$ was only kept if its Euclidean distance from the origin was less than or equal a distance specified by the following equations:

$$\sqrt{x^{2}+ y^{2}} \leq1-2\sin\left( \theta\right)+\cos\left( \theta^{2} \right)$$

$$\theta=\text{atan2}(y,x)$$

*Rasterization*

For the 100 random simulated datasets, we formatted the position and gene expression from each dataset into `SpatialExperiment` objects using the R package *SpatialExperiment* version 1.18.1^6^. These `SpatialExperiment` objects were rasterized into hexagonal pixels with resolution = 0.2 using R package *SEraster* version 1.0.0^7^, such that each pixel value is the mean of gene expression of all simulated cells within the boundaries of the pixel.

*Calculation of Spatial Correlation*

For every pair combination possible for the 100 random simulated datasets except the self-pairs, we used the `spatialCorrelationGeneExp` function to calculate the correlation coefficient, the analytical p-value, and the empirical p-value to compare the rasterized gene expression in the pair. We used the default parameters in which $B=100$ permutations and $\delta= [0.1, 0.2, 0.3, 0.4, 0.5, 0.6, 0.7, 0.8, 0.9]$.

4. MERFISH

*Data Acquisition for MERFISH Brain Data*

Two ST datasets of mouse brain coronal sections assayed by MERFISH were obtained from Clifton et al^9^. We chose Slice 2 Replicate 2 (S2R2) and Slice 2 Replicate 3 (S2R3) as the two biological replicates from matched locations with respect to bregma. The positions of single cells in S2R3 had already been spatially aligned by STalign and by affine alignment to S2R2 as previously described^10^.

*Preprocessing Data*

For MERFISH datasets S2R2 and S2R3, we formatted the position and gene expression from each dataset into `SpatialExperiment` objects using the R package *SpatialExperiment* version 1.18.1^6^. These two `SpatialExperiment` objects were rasterized into 200µm resolution square pixels using R package *SEraster* version 1.0.0^7^ , such that for a gene each pixel value is the mean of gene expression of all cells within the boundaries of the pixel.

*Spatially Variable Gene Analysis*

We identified SVGs using Moran’s I as implemented in R package *MERINGUE* version 1.0^5^. To build the adjacent weight matrix for the MERFISH brain datasets, we used filterDist= 200µm as the distance beyond which two pixels cannot be neighbors. For the implementation of Local Indicators of Spatial Autocorrelation (LISA) which enables the process to filter out spatial patterns driven by small numbers of pixels, we set the minimum percent of pixels or spots that must be driving the spatial pattern to 1%. Once we performed SVG analysis on both rasterized versions of MERFISH brain datasets S2R2 and S2R3, we classified the intersection of the resulting lists as the SVGs in both datasets and the symmetric difference as “non-SVGs”.

*Calculation of Spatial Correlation*

For the rasterized versions of the MERFISH datasets S2R2 and S2R3, we used the `spatialCorrelationGeneExpIterPermutations` function to calculate the correlation coefficient (and the associated empirical p-value) for the rasterized gene expression of every gene. This function takes a list $B=[100, 1000]$ and first uses $B=100$ permutations and then uses $B=1000$ permutations for genes for which the BH-adjusted empirical p-value is < 0.05. We set $\delta= [0.01, 0.05, 0.1, 0.2, 0.3, 0.4, 0.5, 0.6, 0.7, 0.8, 0.9]$.

*Calculation of Spatial Similarity*

For the rasterized versions of MERFISH datasets S2R2 and S2R3, we used the `spatialSimilarity` function to calculate the similarity score for the rasterized expression of every gene using $b=1$ and $q=0.05$.

5. Visium

*Data Acquisition for Visium Kidney Data*

Two ST datasets of mice left kidney sections assayed by 10X Genomics Visium at 55um spots were obtained from the authors of Gharaie et al^11,12^. For the 24-hour after ischemia acute kidney injury (AKI) dataset, we used sample “IL3”. For a control dataset, we used sample “NL3”.

*Joint Clustering of Visium Kidney Datasets into Anatomical Regions*

For both the 24hr AKI and control datasets, we performed log_10_ counts per million (CPM) library normalization with a pseudocount of 1 for each gene and filtered the genes to only include those which were expressed in at least 1% of spots. Then, for the intersection of the genes that satisfy that criteria in both datasets, we combined the data and calculated the first 30 principal components (PC) using principal component analysis as implemented in the R package *irlba^13^*. We harmonized the PCs using `HarmonyMatrix` from R package *harmony* version 1.2.4^14^ increasing the diversity clustering penalty parameter, theta, to 10 to result in more diverse clusters. To verify the success of harmonization, we performed dimensionality reduction on the harmonized PCs using `Rtsne` from the R package *Rtsne* version 0.17^15^ and plotted the resulting 2D embedding to confirm that the batches were well mixed. The harmonized PCs were clustered using Louvain clustering as implemented in MERINGUE^5^. Setting the k-nearest neighbor parameter to k=100 resulted in eight clusters. From visual inspection and knowledge of kidney anatomy, these eight clusters were reduced into four clusters: cortex, outer medulla, inner medulla, and other.

*Alignment and Rasterization*

After labeling the Visium spots, we used STalign to perform an affine-only alignment of the AKI to the control based generating images from one-hot encoding of region labels: cortex, outer medulla, and inner medulla^12^. We formatted the aligned positions and gene expression from each dataset into `SpatialExperiment` objects using the R package *SpatialExperiment* version 1.18.1^6^. These two `SpatialExperiment` objects were rasterized into 5 unit resolution hexagonal pixels using R package *SEraster* version 1.0.0^7^ , such that for a gene each pixel value is the sum of gene expression of all cells within the boundaries of the pixel. Following rasterization, we performed CPM library normalization for each gene.

*Spatially Variable Gene Analysis*

We identified SVGs using Moran’s I as implemented in R package *MERINGUE* version 1.0^5^. To build the adjacency weight matrix for the rasterized Visium kidney datasets, we set filterDist = 10 so only the most adjacent spots are neighbors. For the Visium AKI and control datasets, after we performed SVG analysis on each, then we only performed future analysis on the genes that were SVGs in both datasets.

*Calculation of Spatial Correlation*

For the Visium AKI and control datasets, we used the `spatialCorrelationGeneExpIterPermutations` function to calculate the correlation coefficient (and the associated BH-adjusted empirical p-value) for the rasterized library normalized expression of every gene. We used the parameters and $\delta= \left[ 0.01, 0.05, 0.1, 0.2, 0.3, 0.4, 0.5, 0.6, 0.7, 0.8, 0.9 \right]$ and $B=[100, 1000]$ permutations.

*Calculation of Spatial Similarity*

For the Visium AKI and control datasets, we used the `spatialSimilarity` function to calculate the similarity score for the library normalized expression of every gene using $b=1$ and $q=0.05$.

6. Comparison with Other Methods

*STdiff*: *Data Preprocessing*

The authors implement `STdiff` as a function in the `spatialGE` package. As `STdiff` was designed for domain-specific differential expression testing within a sample, the function expects one dataset with domain annotations for each cell or spot. To use `STdiff` for differential expression testing across paired samples, we combined our gene expression pattern pairs and provided domain annotations that map to each gene expression pattern. For example, gene expression patterns A and B are joined such that gene expression pattern A is domain 1 and gene expression pattern B is domain 2.

Furthermore, the authors’ implementation of `STdiff` does not enable differential expression testing on a dataset with only one gene. As testing spatial models with `STdiff` is computationally intensive with respect to both memory and time, the `STdiff` function tries to reduce the number of genes in the dataset by enforcing a calculation to get the most highly variable genes but the implementation results in an error that terminates the function if there are not enough genes to perform the calculation correctly. To avoid this error, we supplement the simulated gene in gene expression patterns A and B with 99 additional simulated genes. We simulated the expression of these additional genes to be less variable than the original gene. The expression of each additional gene is from random sampling from a normal distribution with μ = mean of the original simulated gene and σ=1/2*standard deviation of original simulated gene.

*STdiff*: *Differential Expression Testing*

We used the `STlist` function from `spatialGE` to coerce our `SpatialExperiment` objects into `STlist` objects. Then, rather than using the `STclust` function to generate the domain annotations, we manually added the appropriate domain annotations to the `spatial_meta` feature of the `STlist` object. Finally, we set the `STdiff` function parameters to `topgenes = 1` and `sp_topgenes=1` to only perform differential expression tests on the single most highly variable gene.

*SPADE*: *Data Preprocessing*

Since dataset pairs A, B, and C only have one gene each we did not use the `SPADE_norm` function which performs library size normalization.

*SPADE*: *Spatially Variable Gene Analysis*

We used the default parameters for the `SPADE_DE` function for both gene expression pattern pair A and B and gene expression pattern pair A and C.

*SPaSE*: *Data Preprocessing*

To use SPaSE we converted the rasterized, CPM normalized AKI and control Visium SpatialExperiment objects into AnnData objects.

SPaSE:

We used `run.py` to execute SPaSE with the file path to the control `.h5ad` as the left (healthy) path and the file path to the AKI `.h5ad` as the right (diseased) path. We set the parameters as alpha = 0.01 and lambda_sinkhorn = 0.1. Additionally, in order for SPaSE to create the synthesized null data from the control as described by the authors of SPaSE we corrected the `AnalyzeOutput.py`to uncomment line 156.

7. Runtime Benchmarking

To evaluate the runtime scalability of spatial correlation test, we benchmarked runtime across datasets containing varying numbers of spatial locations. Benchmarks were run on a Mac Studio with an Apple M2 Ultra processor, 24 CPU cores (16 performance and 8 efficiency cores), and 192 GB RAM, running macOS 15.6.1 and R 4.3.3. The spatial correlation procedure was run with 22 threads using BiocParallel, and runtime was measured for the spatial correlation test.

Runtime benchmark datasets were generated by rasterizing the two biological replicates of coronal slices of the adult mouse brain from different animals but the same location with respect to bregma assayed by the single-cell resolution spatial transcriptomics technology MERFISH (Figure 2a) at resolutions of 50, 100, 150, 200, 250, and 300, resulting in 176, 367, 598, 839, 1088, and 1371 spots, respectively. For each resolution, the spatial correlation test was applied to the entire gene panel, 483 genes.

For each dataset size, spatialCorrelationGeneExpIterPermutations() was run five times using different random seeds to capture variation arising from the permutation step. Total wall-clock runtime was recorded for each run and normalized by the number of genes analyzed to obtain the mean runtime per gene. For each dataset size, the mean and standard deviation of runtime per gene were calculated across replicate runs. To quantify runtime scaling, a linear regression model was fit to the relationship between the number of spatial locations and mean runtime per gene. The slope of the fitted regression line was used to estimate the increase in runtime cost associated with each additional spatial location.

**Supplementary Figures**

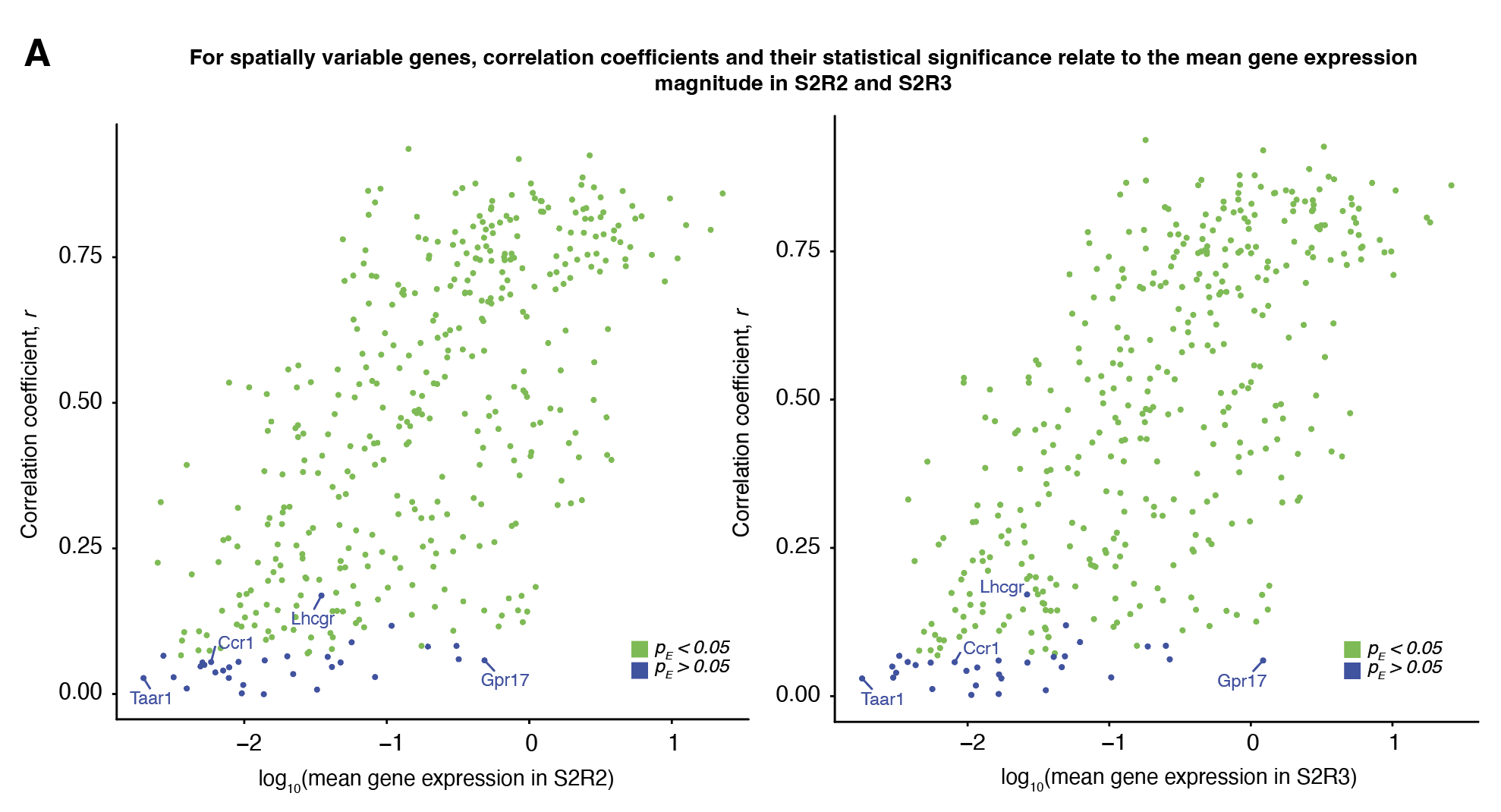

**Supplementary Figure 1. *Spatially variable genes with non-significant spatial correlation across mouse brain coronal slice replicate have low expression.* A**. For each of the 415 genes that are SVG in both Slice 2 Replicate 2 (S2R2) and Slice 2 Replicate 3 (S2R3), we calculate the correlation coefficient from comparing S2R2 and S2R3 at matched locations and compare the coefficient to the log_10_ mean expression of the gene across all pixels in S2R2 (left) and S2R3 (right). Genes with non-significant spatial correlation, empirical p-value > 0.05 (blue), and genes with have significant spatial correlation, empirical p-value < 0.05 (green).

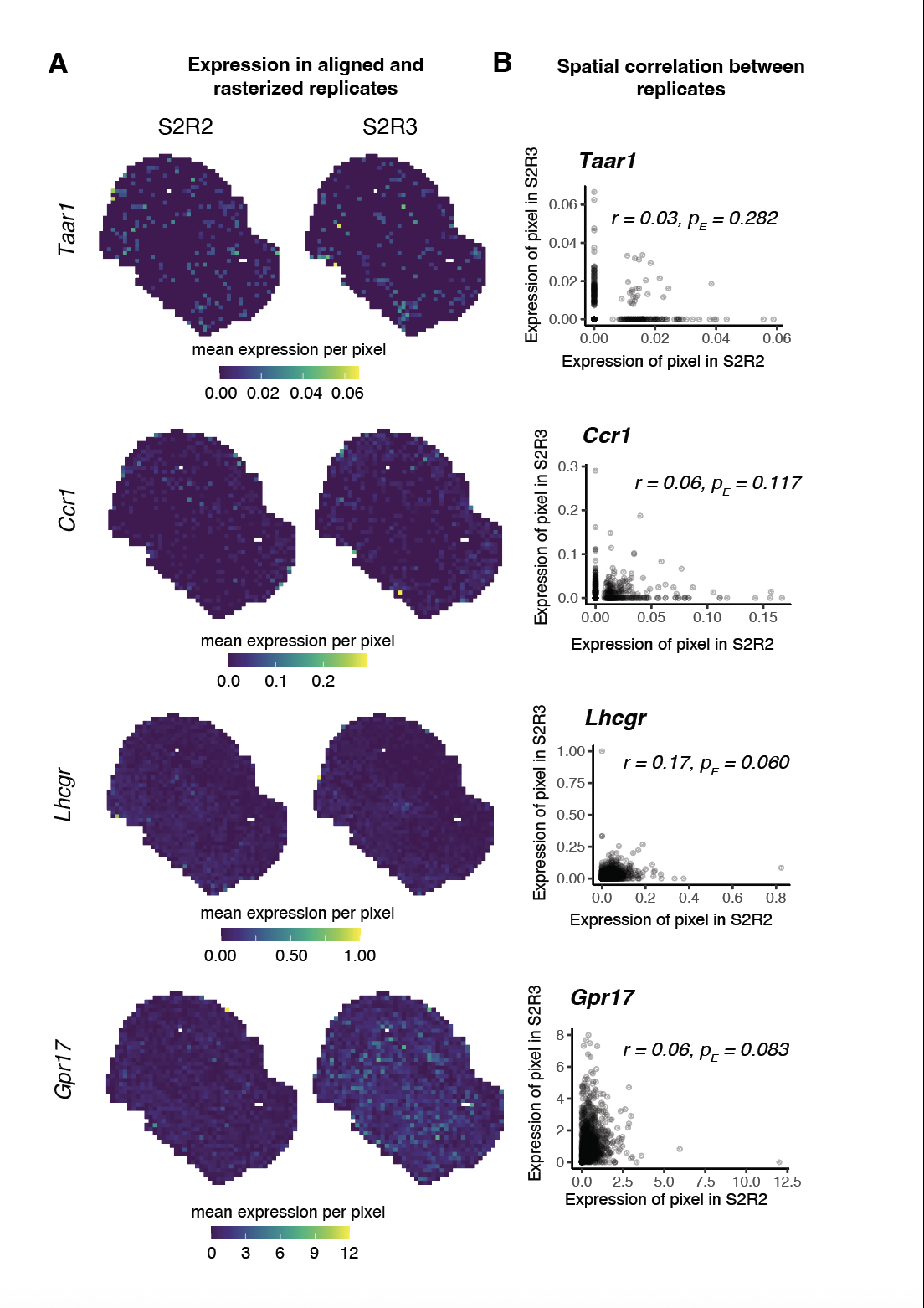

**Supplementary Figure 2. *Examples of spatially variable genes with low and non-significant spatial correlation across mouse brain coronal slice replicates.* A**. For examples of spatially variable genes that are not significantly positively correlated, visualizations of the mean expression per pixel in the aligned and rasterized replicates S2R2 (left) and S2R3 (right). B. For these select genes, plots of expression in S2R2 versus S2R3 at matched pixels with correlation coefficient and empirical p-value.

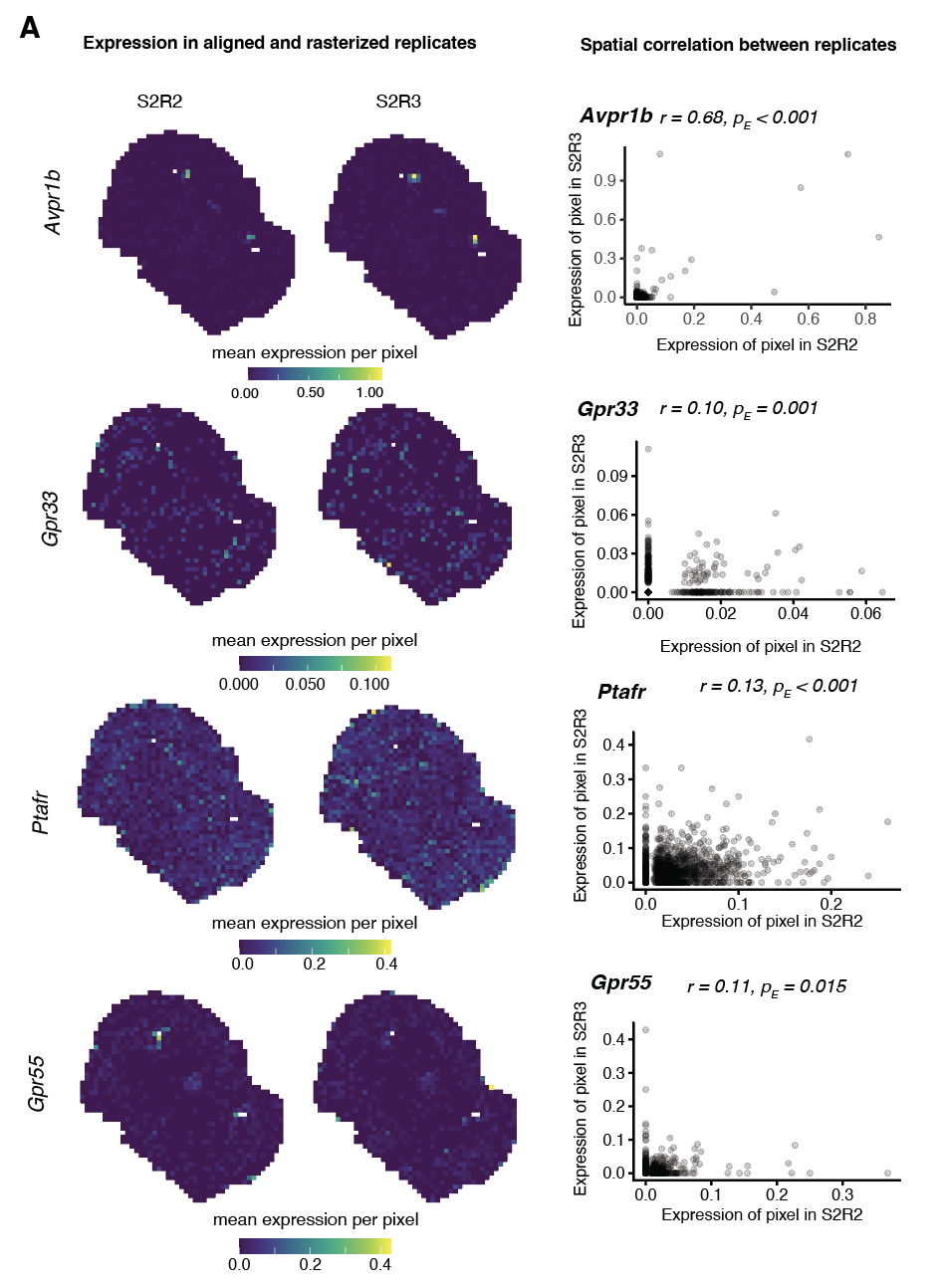

**Supplementary Figure 3. Examples of *non-spatially variable genes with significant positive spatial correlation (r > 0, p_E_ < 0.05) across mouse brain coronal slice replicates.*** **A.** Visualizations of the mean expression per pixel in the aligned and rasterized replicates, S2R2 (left) and S2R3 (middle) for examples of not spatially variable genes that are significantly positively correlated (right) show that *Avpr1b* has clustered patterns with area smaller than 1% of pixels, *Gpr33* has patterns that are organized but with high dispersion, and *Ptafr* and *Gpr55* have low expression with outliers could have impacted whether the genes are classified as SVG as in both datasets.

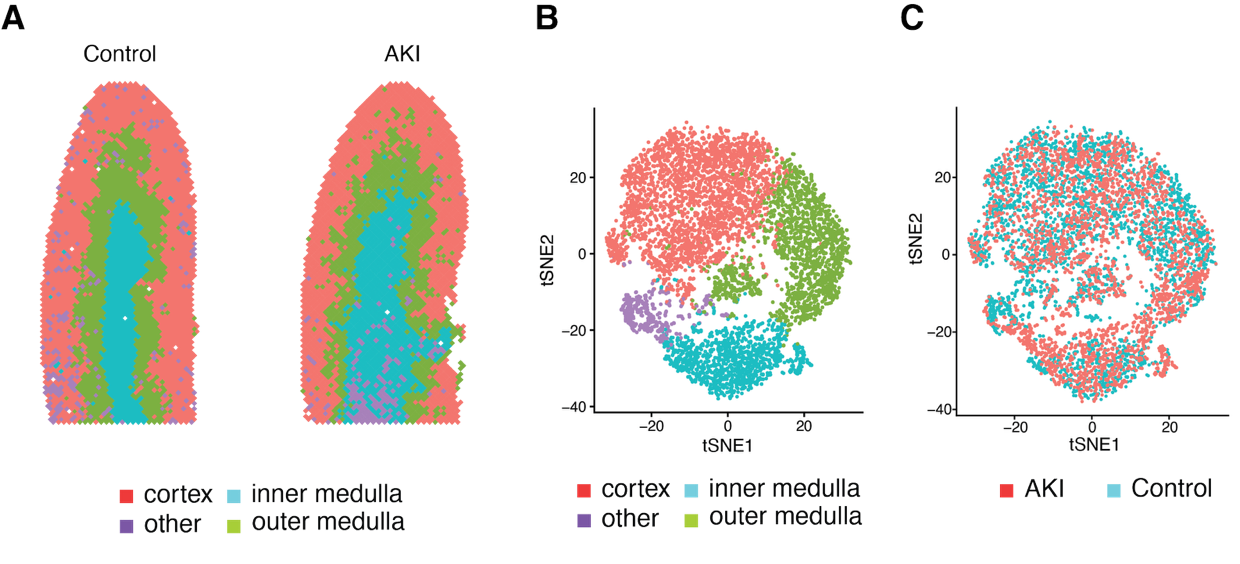

**Supplementary Figure 4. Harmonized transcriptional clustering of spots from two kidney sections, control and AKI, assayed by 10x Visium***.* **A**. kidney compartments in control and AKI as annotated via harmonized Louvain clustering **B**. tSNE with harmonized Louvain clusters annotated as kidney compartments. **C**. tSNE with harmonized Louvain clusters annotated with sample ID of control or AKI shows the batches are well mixed with each cluster including spots from both tissue sections.

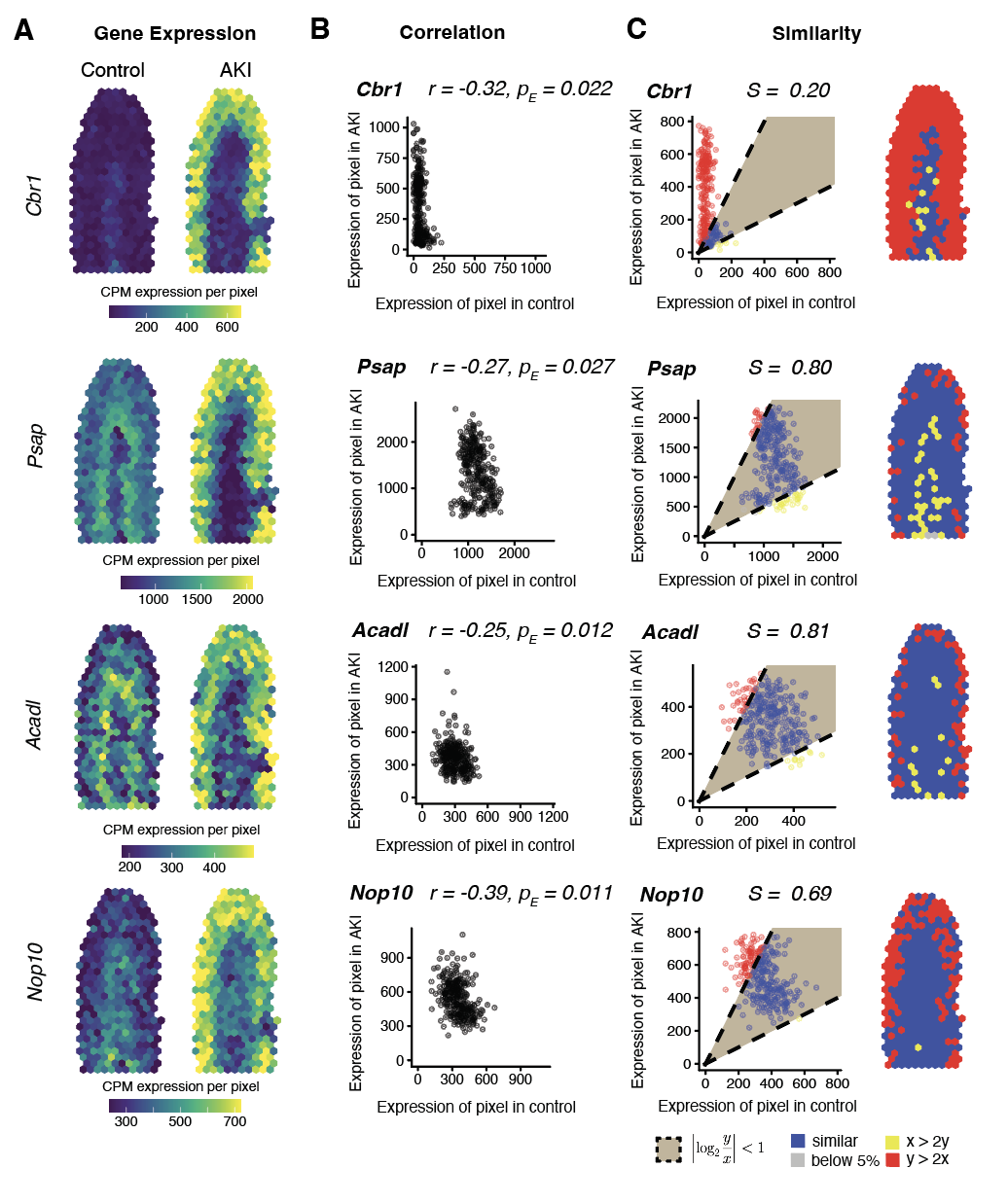

**Supplementary Figure 5. *Examples of significantly spatially differential genes (r < 0, p_E_ < 0.05) with decreased expression in medulla and increased expression in cortex in ischemia compared to control***

**A.** Spatial visualizations of counts per million (CPM) normalized gene expression per spot for control (left) and AKI (right) for examples of significantly positively correlated genes with relatively higher similarity scores **B.** Plot of CPM normalized expression in control versus with AKI at matched spots with correlation coefficient and empirical p-value **C.** Plot (left) of CPM normalized expression in control versus AKI at matched spots with spots classified as either having twice as much expression in AKI (red), having twice as much expression in control (yellow), or similar (blue), and spatial visualization (right) of spot classification. The domain of plots restricted to the 95^th^ quantile of spot based expression.

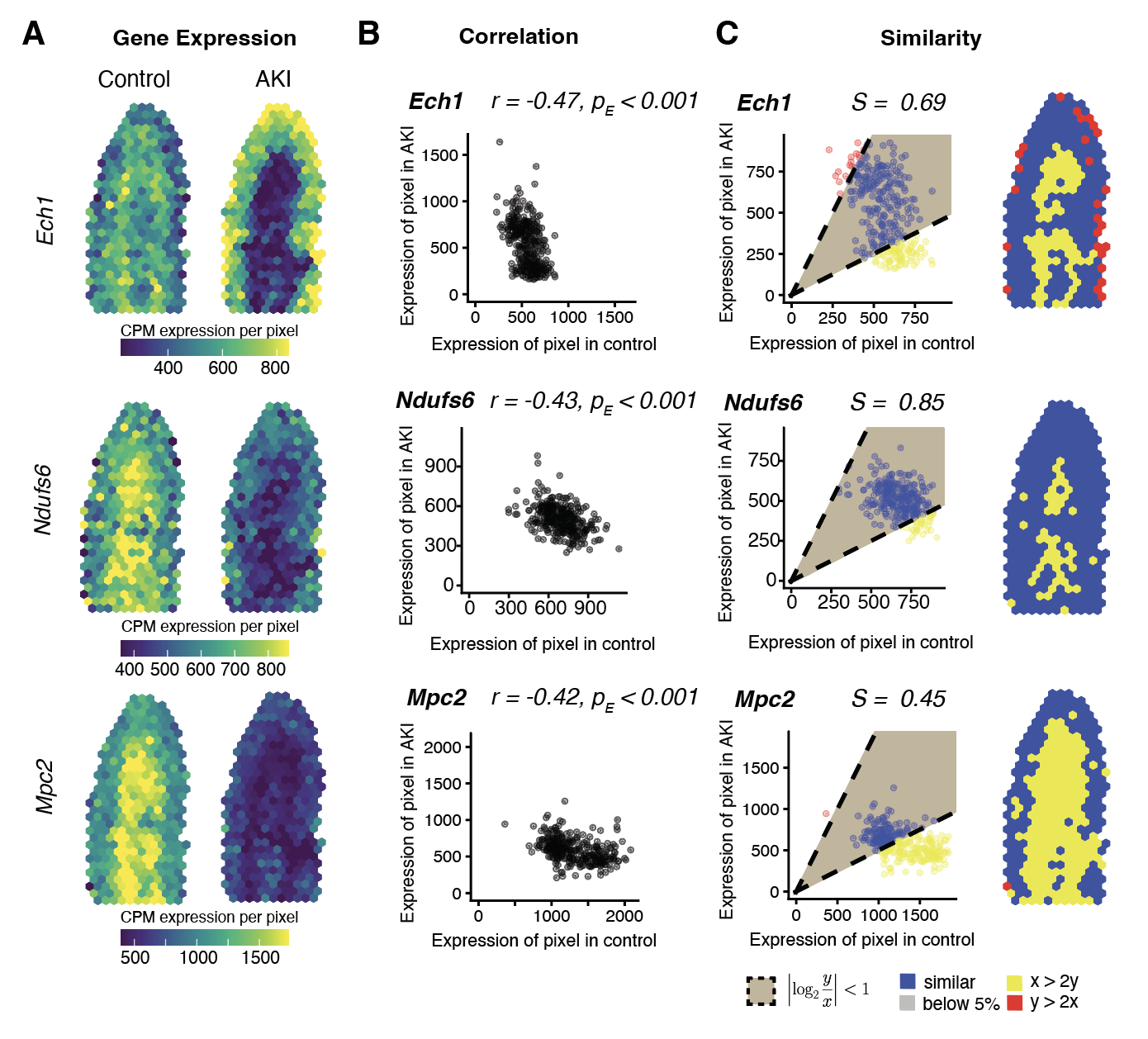

**Supplementary Figure 6. *Examples of significantly spatially differential genes (r < 0, p_E_ < 0.05) with decreased expression in medulla and comparable expression in cortex in ischemia compared to control***

**A.** Spatial visualizations of counts per million (CPM) normalized gene expression per spot for control (left) and AKI (right) for examples of significantly positively correlated genes with relatively higher similarity scores **B.** Plot of CPM normalized expression in control versus with AKI at matched spots with correlation coefficient and empirical p-value **C.** Plot (left) of CPM normalized expression in control versus AKI at matched spots with spots classified as either having twice as much expression in AKI (red), having twice as much expression in control (yellow), or similar (blue), and spatial visualization (right) of spot classification. The domain of plots restricted to the 95^th^ quantile of spot based expression.

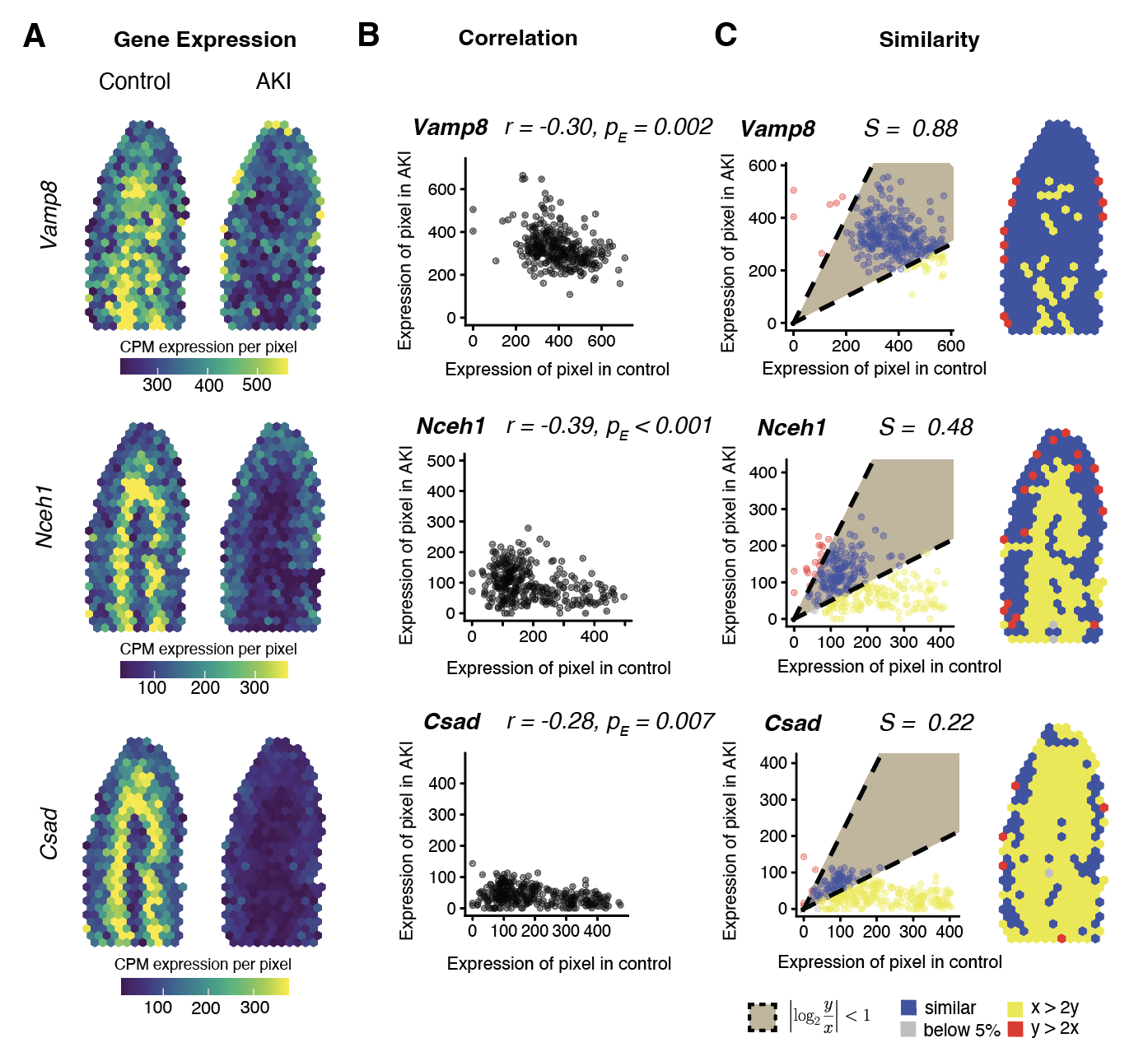

**Supplementary Figure 7. *More examples of significantly spatially differential genes (r < 0, pE < 0.05) with decreased expression in medulla and comparable expression in cortex in ischemia compared to control* A.** Spatial visualizations of counts per million (CPM) normalized gene expression per spot for control (left) and AKI (right) for examples of significantly positively correlated genes with relatively higher similarity scores **B.** Plot of CPM normalized expression in control versus with AKI at matched spots with correlation coefficient and empirical p-value **C.** Plot (left) of CPM normalized expression in control versus AKI at matched spots with spots classified as either having twice as much expression in AKI (red), having twice as much expression in control (yellow), or similar (blue), and spatial visualization (right) of spot classification. The domain of plots restricted to the 95^th^ quantile of spot based expression.

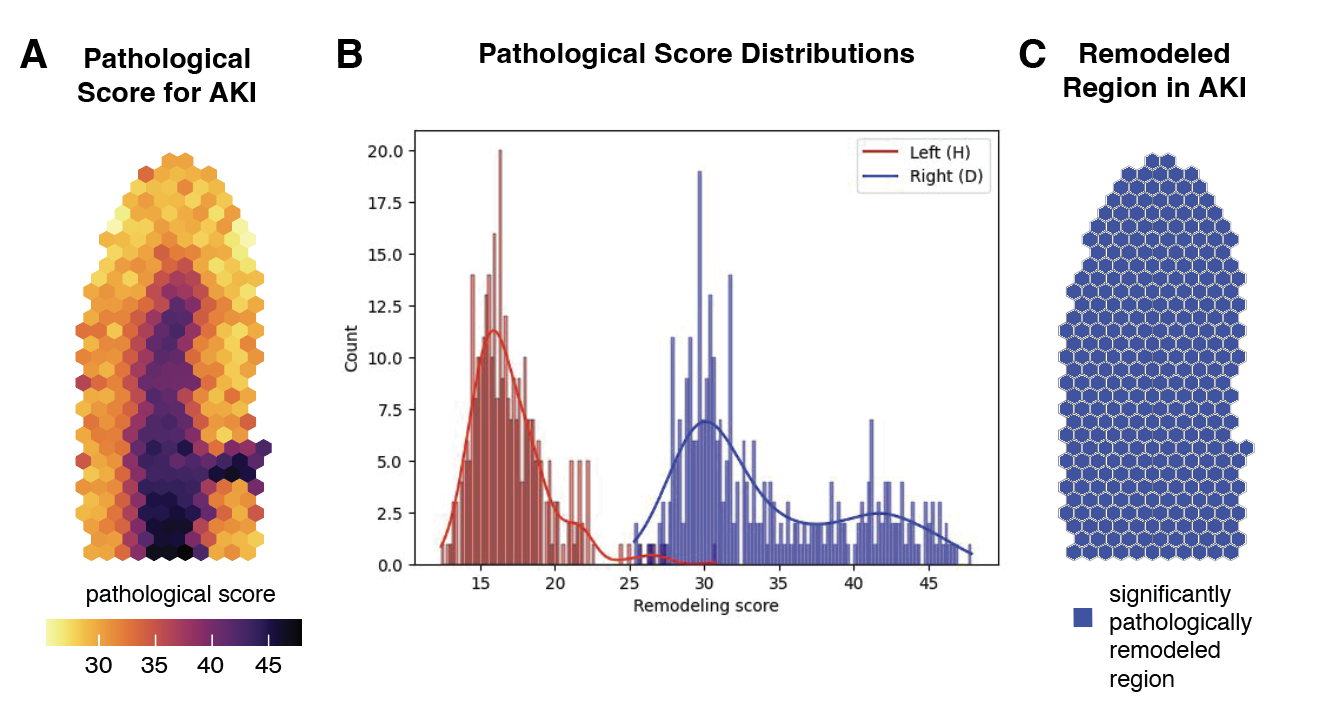

**Supplementary Figure 8*. SPaSE calculates pathological scores and identifies significantly pathologically remodeled region*** **A**. Spatial plot of SPaSE pathological scores calculated for each pixel of rasterized AKI Visium. **B**. Remodeling (aka pathological) scores for each pixel of the rasterized AKI (blue) and control (red) tissues. **C**. Spatial plot of significantly pathologically remodeled region (blue) of rasterized AKI Visium.

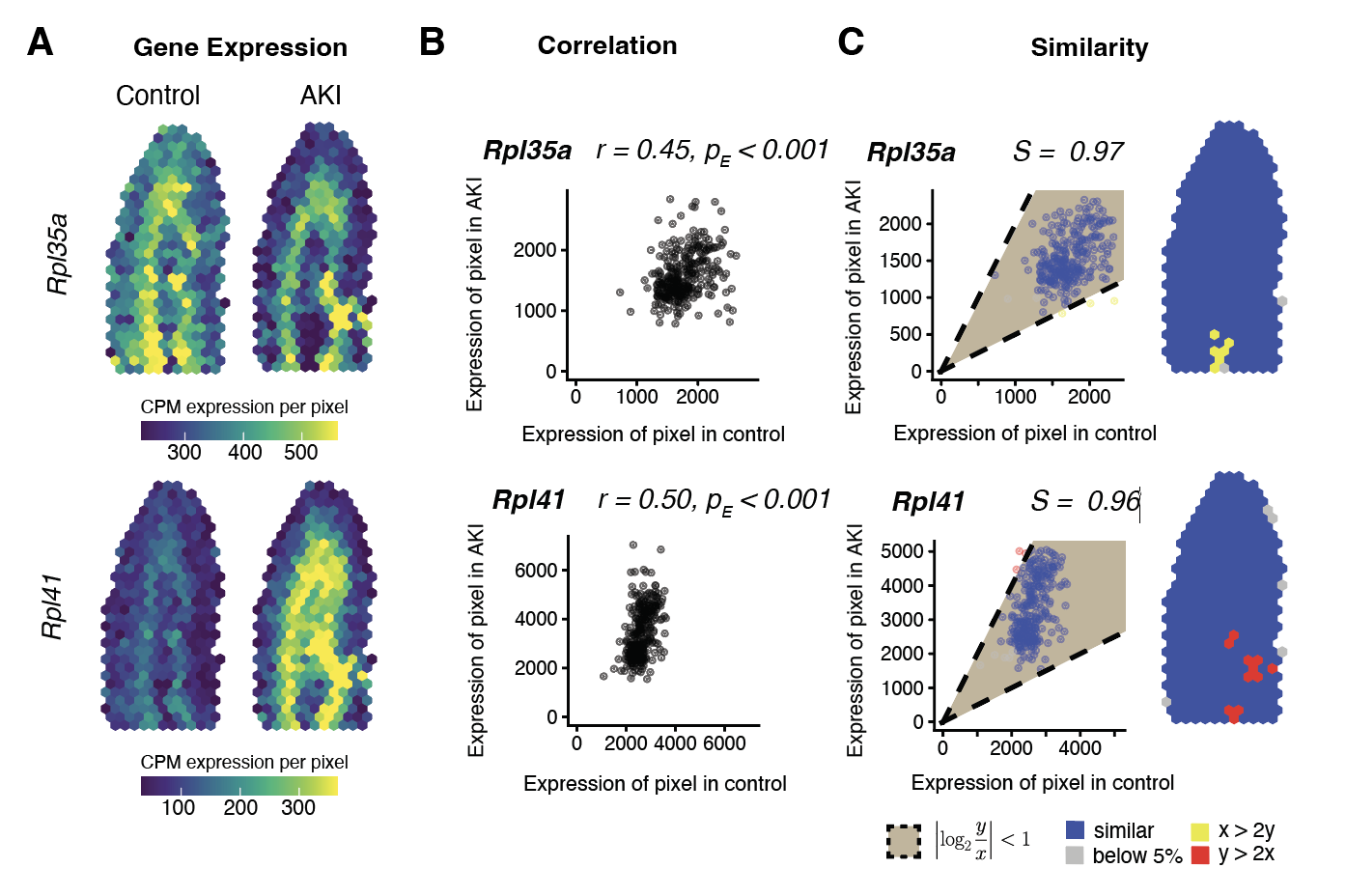

**Supplementary Figure 9. *Examples of significantly similarly spatially patterned genes (r > 0, p_E_ < 0.05) with high similarity scores due to lack of change in expression magnitude*** **A**. Spatial visualizations of counts per million (CPM) normalized gene expression per spot for control (left) and AKI (right) for examples of significantly positively correlated genes with relatively higher similarity scores. **B.** Plot of CPM normalized expression in control versus with AKI at matched spots with correlation coefficient and empirical p-value **C.** Plot (left) of CPM normalized expression in control versus AKI at matched spots with spots classified as either having twice as much expression in AKI (red), having twice as much expression in control (yellow), or similar (blue), and spatial visualization (right) of spot classification. The domain of plots restricted to the 95^th^ quantile of spot based expression.

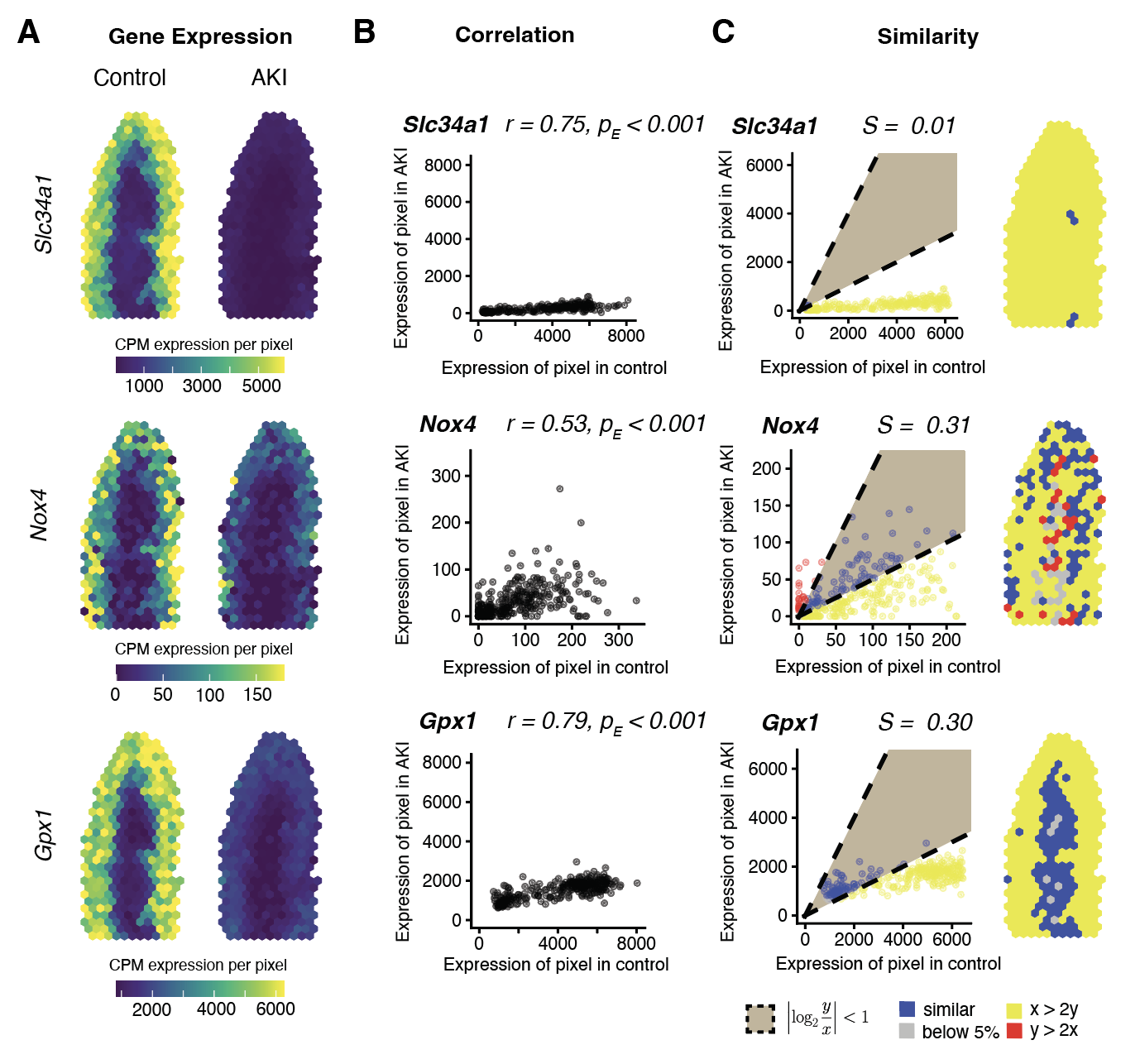

**Supplementary Figure 10. *Examples of significantly similarly spatially patterned genes (r > 0, p_E_ < 0.05) with low similarity scores due to decrease in expression magnitude*** **A.** Spatial visualizations of counts per million (CPM) normalized gene expression per spot for control (left) and AKI (right) for examples of significantly positively correlated genes with relatively higher similarity scores **B.** Plots of CPM normalized expression in control versus with AKI at matched spots with correlation coefficient and empirical p-value **C.** Plots (left) of CPM normalized expression in control versus AKI at matched spots with spots classified as either having twice as much expression in AKI (red), having twice as much expression in control (yellow), or similar (blue), and spatial visualization (right) of spot classification. The domain of plots restricted to the 95^th^ quantile of spot based expression.

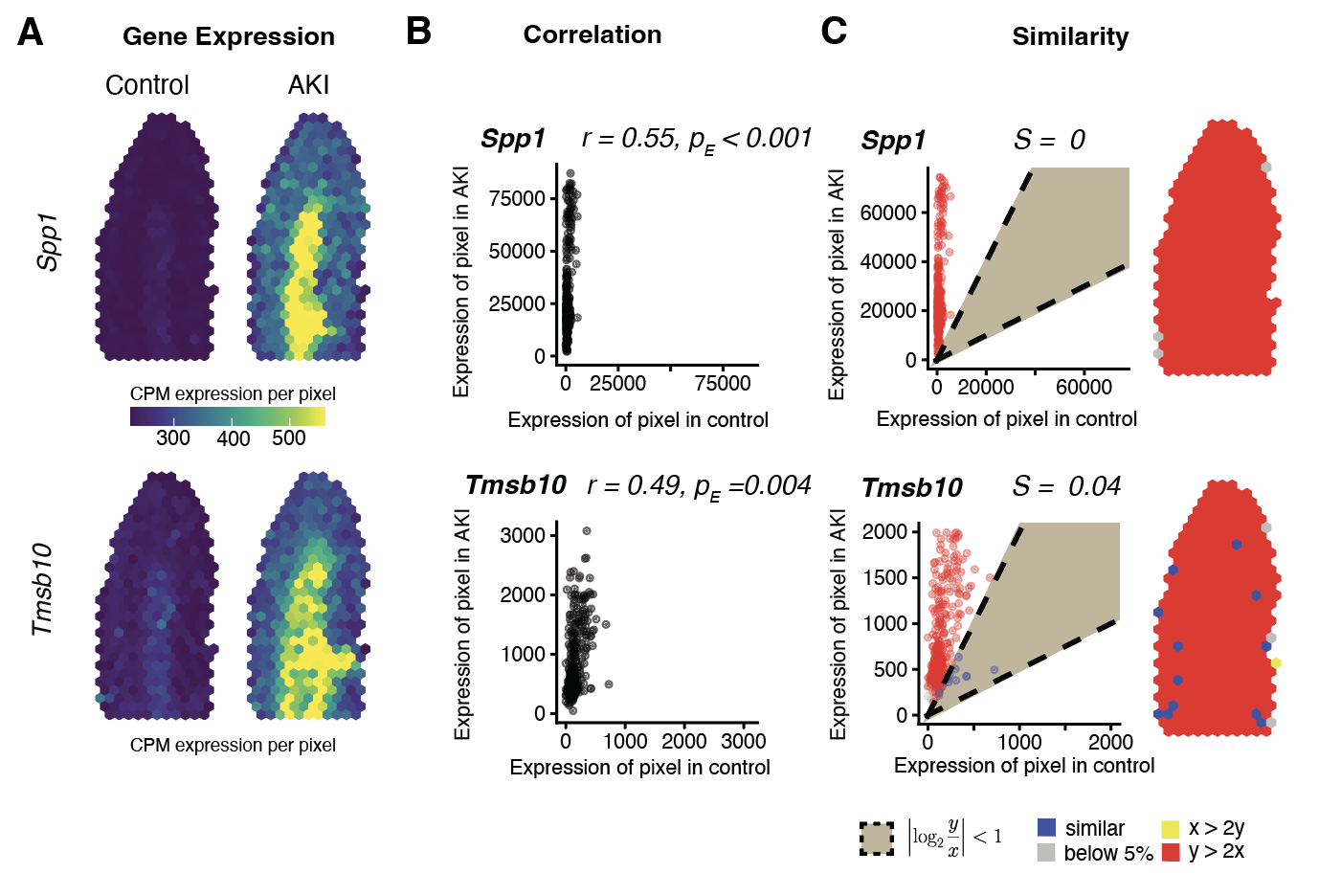

**Supplementary Figure 11. *Examples of significantly similarly spatially patterned genes (r > 0, p_E_ < 0.05) with low similarity scores due to increase in expression magnitude*** **A.** Spatial visualizations of counts per million (CPM) normalized gene expression per spot for control (left) and AKI (right) for examples of significantly positively correlated genes with relatively higher similarity scores **B.** Plots of CPM normalized expression in control versus with AKI at matched spots with correlation coefficient and empirical p-value **C.** Plots (left) of CPM normalized expression in control versus AKI at matched spots with spots classified as either having twice as much expression in AKI (red), having twice as much expression in control (yellow), or similar (blue), and spatial visualization (right) of spot classification. The domain of plots restricted to the 95^th^ quantile of spot based expression.

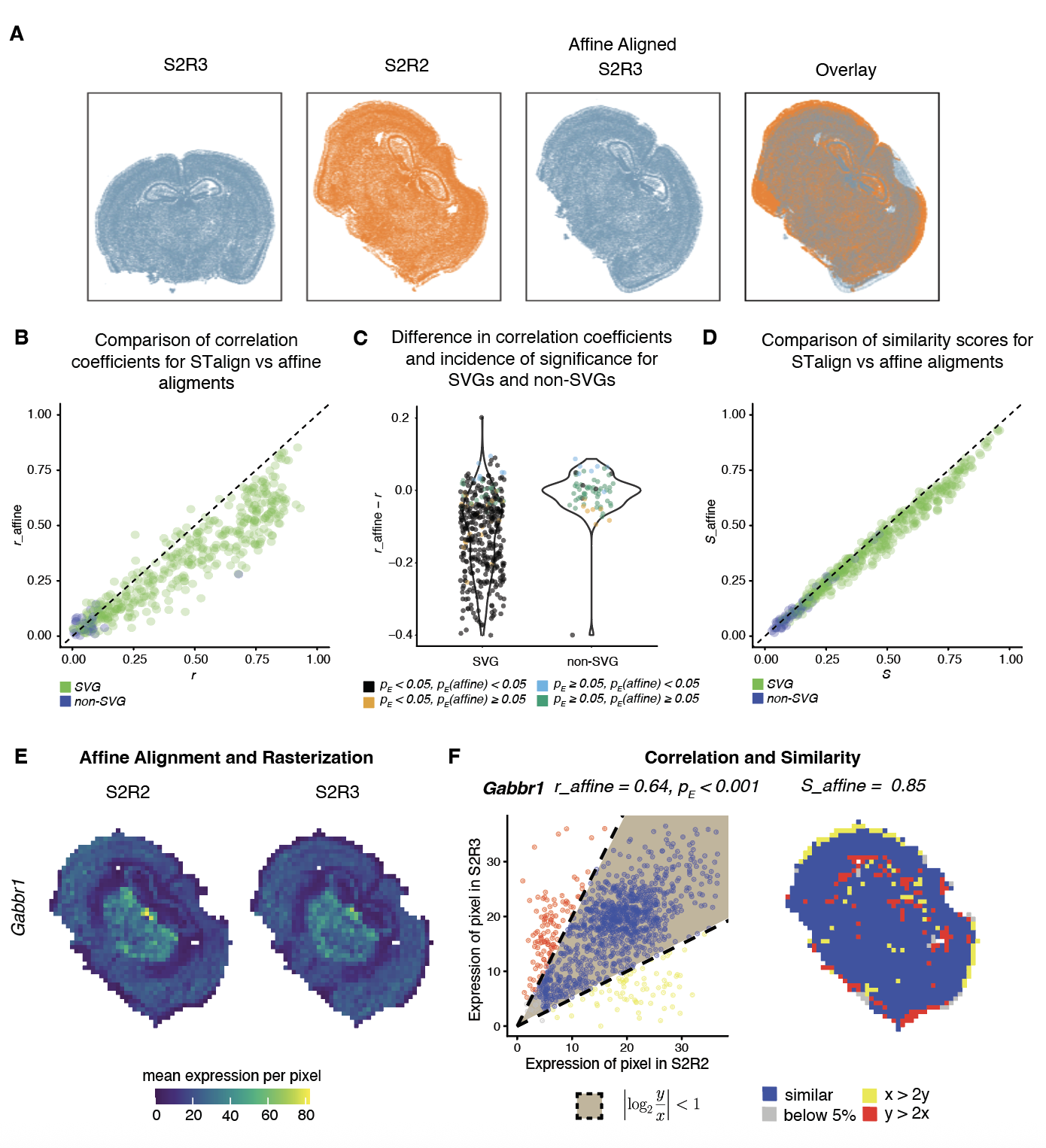

**Supplementary Figure 12. *Evaluation of STcompare with MERFISH datasets aligned with affine alignment.*** **A**. Spatial agreement of S2R3 and S2R2 that has been aligned based on a simple affine transformation based on manually placed landmarks as described in Clifton et al. **B**. Correlation coefficients for S2R2 versus S2R3 for STalign compared to affine alignment for 415 SVGs (green) and 68 non-SVGs (blue). **C**. Difference in correlation coefficients for alignment methods (r_affine – r) for 415 SVGs and 68 non-SVGs. Each gene is colored regarding with which alignment method it was identified as statistically significantly correlated: with both STalign and affine (black), with STalign but not with affine (orange), with affine but not STalign (blue), and with neither STalign nor affine (green). **D**. Similarity scores for S2R2 versus S2R3 for STalign compared to affine alignment for 415 SVGs (green) and 68 non-SVGs (blue). **E**. Visualizations of the mean expression of *Gabbr1* per pixel in S2R2 (left) and S2R3 (right) after affine alignment and rasterization. **F**. Plot (left) of expression in S2R2 versus S2R3 at matched pixels with correlation coefficient and empirical p-value for *Gabbr1,* r_affine = 0.64, p_E_ < 0.001. Pixels classified as either having twice as much expression in S2R3 (red), having twice as much expression in S2R2 (yellow), or similar expression (blue). Spatial visualization (right) of spot classification with similarity scores for *Gabbr1* (S_affine = 0.85). Plot domains restricted to the 95^th^ quantile of pixels-based expression.

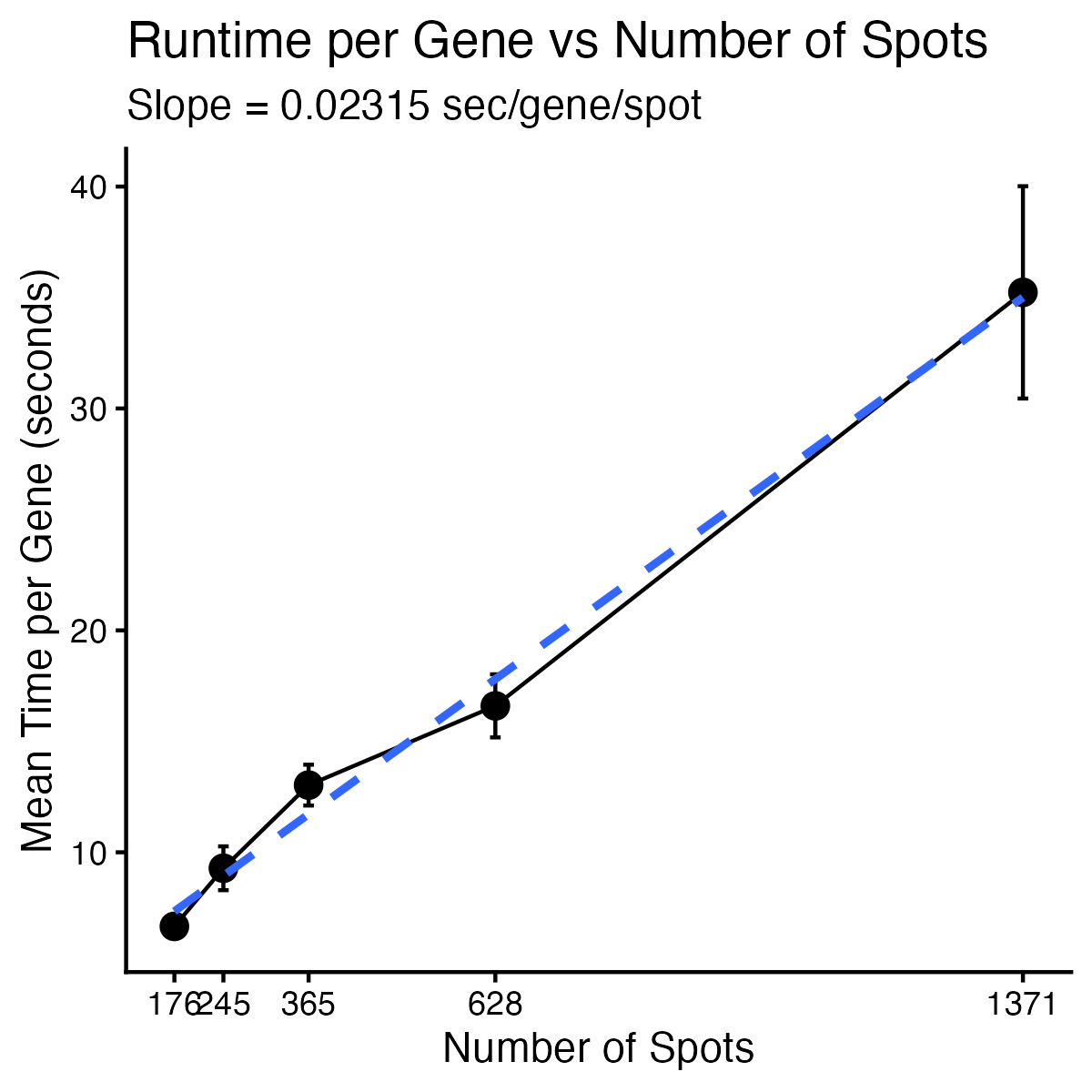

**Supplementary Figure 13. Runtime scaling of STcompare spatial correlation analysis.** Mean runtime per gene as a function of the number of spatial locations (spots). Points represent the mean across benchmarking runs and error bars indicate one standard deviation. The dashed line shows a linear regression fit.

Supplementary References

1.	R Core Team. R: A Language and Environment for Statistical Computing. Preprint at https://www.R-project.org/ (2025).

2.	Viladomat, J., Mazumder, R., Mcinturff, A., Mccauley, D. J. & Hastie, T. Assessing the Significance of Global and Local Correlations under Spatial Autocorrelation: A Nonparametric Approach. *Biometrics* **70**, 409–418 (2014).

3.	Ribeiro Jr, P. J. & Diggle, P. geoR: Analysis of Geostatistical Data. *CRAN: Contributed Packages* Preprint at https://doi.org/10.32614/CRAN.package.geoR (2001).

4.	Loader, C., Sun, J., Lucent Technologies & Liaw, A. locfit: Local Regression, Likelihood and Density Estimation. *CRAN: Contributed Packages* Preprint at https://doi.org/10.32614/CRAN.package.locfit (2025).

5.	Miller, B. F., Bambah-Mukku, D., Dulac, C., Zhuang, X. & Fan, J. Characterizing spatial gene expression heterogeneity in spatially resolved single-cell transcriptomic data with nonuniform cellular densities. *Genome Res.* **31**, 1843–1855 (2021).

6.	Righelli, D. *et al.* SpatialExperiment: infrastructure for spatially-resolved transcriptomics data in R using Bioconductor. *Bioinformatics* **38**, 3128–3131 (2022).

7.	Aihara, G. *et al.* SEraster: a rasterization preprocessing framework for scalable spatial omics data analysis. *Bioinformatics* **40**, (2024).

8.	Venables, W. N. & Ripley, B. D. *Modern Applied Statistics with S*. (Springer, New York, 2002).

9.	Clifton, K., Anant, M. & Fan, J. STalign: Alignment of spatial transcriptomics data using diffeomorphic metric mapping [Data set]. *Zenodo* Preprint at (2023).

10.	Clifton, K. *et al.* STalign: Alignment of spatial transcriptomics data using diffeomorphic metric mapping. *Nat. Commun.* **14**, 8123 (2023).

11.	Gharaie, S. *et al.* Single cell and spatial transcriptomics analysis of kidney double negative T lymphocytes in normal and ischemic mouse kidneys. *Sci. Rep.* **13**, 20888 (2023).

12.	Clifton, K., Fan, J. & Rabb, H. STcompare: comparative spatial transcriptomics data analysis of structurally matched tissues to characterize differentially spatially patterned genes [Data set]. *Zenodo* Preprint at https://doi.org/https://doi.org/10.5281/zenodo.20647680 (2026).

13.	Baglama, J., Reichel, L. & Lewis, B. W. irlba: Fast Truncated Singular Value Decomposition and Principal Components Analysis for Large Dense and Sparse Matrices. *CRAN: Contributed Packages* Preprint at https://doi.org/10.32614/CRAN.package.irlba (2011).

14.	Korsunsky, I. *et al.* harmony: Fast, Sensitive, and Accurate Integration of Single Cell Data. *CRAN: Contributed Packages* Preprint at https://doi.org/10.32614/CRAN.package.harmony (2025).

15.	Krijthe, J. & van der Maaten, L. Rtsne: T-Distributed Stochastic Neighbor Embedding using a Barnes-Hut Implementation. *CRAN: Contributed Packages* Preprint at https://doi.org/10.32614/CRAN.package.Rtsne (2023).
